# Self-generated environmental feedback drives traveling waves of gene expression in bacterial colonies

**DOI:** 10.64898/2026.06.16.732571

**Authors:** Guillermo Yáñez Feliú, Conor Brownson, Timothy J. Rudge

**Affiliations:** School of Computing, Newcastle University, Newcastle upon Tyne, NE1 7RU, United Kingdom

**Keywords:** bacterial colonies, nonlinear dynamics, traveling waves, gene expression, envi-ronmental feedback, synthetic biology

## Abstract

Bacterial colonies grow within microenvironments that they continuously reshape through nutrient uptake, metabolism and mechanical interaction. However, in colonies carrying engineered gene circuits, how these self-generated environmental changes feed back on gene expression to produce spatiotemporal organization remains poorly understood. Here we show that growth and gene expression are dynamically coupled during the maturation of founding colonies, with growth-driven environmental changes organizing gene expression into traveling waves. By combining quantitative time-lapse microscopy, mathematical modeling and image-based parameter inference, we demonstrate that edge-dominated colony expansion consistent with mechanical constraints on growth is followed by density-dependent growth arrest in which the total area occupied by colonies does not converge to a fixed carrying capacity of the shared growth environment. We then find that colonies form two distinct traveling waves of gene expression. An intra-colony wave emerges only after growth arrest and is observed in both constitutive genes and distinct regulated circuit architectures, indicating that it is not a result of circuit topology. Our observations are consistent with nutrient depletion during growth followed by subsequent recovery after growth arrest. At later times, an inter-colony wave emerges that is consistent with a colony-produced diffusible factor spreading through the shared medium. Together, these findings reveal that the colony environment is not a passive background but an active and intrinsic component of spatiotemporal gene regulatory dynamics, in which self-generated environmental feedback couples mechanical constraints, nutrient dynamics and gene expression across spatial scales.

## 1 Introduction

Multicellularity and collective cell behavior represent a major evolutionary transition in which individual cells come together to form organized structures with emergent properties [1, 2]. The self-organized spatiotemporal patterns that arise during development, such as traveling waves, differentiated domains, and synchronized oscillations, exemplify how local cellular interactions can generate global order without centralized control [3–5]. Bacterial biofilms, communities of cells attached to a surface and coordinating their physiology through shared signals and physical contact, provide a powerful and tractable model for studying these principles [6, 7]. They are also well suited for studying nonlinear dynamics in living systems, because local interactions between growth, mechanics, metabolism and gene regulation can generate collective spatiotemporal patterns [8–11].

Biofilm formation involves the emergence of spatially organized patterns of gene expression and growth, driven by mechanical forces, nutrient gradients, and intercellular signaling [9, 11–15]. At the colony scale, cells at the periphery grow faster than those in the interior consistent with mechanical constraints from cell-cell and cell-substrate interactions, generating a characteristic gradient of growth rate that decays exponentially from the edge inward [16–19]. Critically, growing colonies do not simply respond to a fixed environment. Cell growth modifies local mechanical and chemical conditions, depleting nutrients, altering density, and changing the physical properties of the shared medium, which can in turn feed back on gene expression [11, 15, 20, 21]. Related self-generated mechanisms have been described in eukaryotic collective migration and vertebrate development, including cell-generated chemoattractant gradients and mechanochemical feedback between tissue rigidity and morphogen transport [22–24]. Because colonies develop within a shared environment, such local dynamics may also couple across colonies, giving rise to organization at the community scale [25].

Synthetic biology has provided a powerful approach to studying these principles [26–28]. By engineering gene circuits with defined architectures into bacterial colonies, one can ask which aspects of observed patterning are intrinsic to the circuit and which arise from the physical and chemical environment in which the circuit operates [29–32]. However, in colonies carrying engineered genetic constructs, it remains unclear how the environment generated by colony growth feeds back on gene expression to produce spatiotemporal organization. The simplest genetic architecture for testing these dynamics is constitutive fluorescent reporter expression, which provides a readout of how colony physiology and environmental conditions shape gene expression in the absence of engineered regulatory interactions [33].

A more complex circuit is the repressilator - a synthetic three-node ring oscillator first implemented in *E. coli* by Elowitz and Leibler [34] and significantly improved by Potvin-Trottier et al. [35]. This system forms concentric ring patterns of gene expression in growing colonies [35], but how colony growth, metabolism, and local environmental conditions shape these patterns remains to be determined. Mathematical modeling predicts that growth rate decays exponentially from the colony edge inward, generating a characteristic radial velocity profile, and suggests that the coupling of gene expression to this growth gradient could in principle generate emergent spatiotemporal patterns in these colonies [19]. More generally, the behavior of engineered gene circuits during colony development remains difficult to predict because growth continuously reshapes the mechanical and chemical environment in which those circuits operate [11, 33].

Here we use engineered gene circuits as quantitative reporters and perturbable architectures to show that the interplay of cell mechanics, environmental conditions, and cell metabolism can generate traveling waves of gene expression across and between colonies. We followed *Escherichia coli* colonies carrying various gene circuits continuously from single cells to colonies exceeding one millimeter in diameter using quantitative time-lapse fluorescence microscopy and applied a custom image-based analysis framework to the resulting data.

We show that (i) colony expansion is edge-dominated, consistent with mechanical constraint predictions [19]; (ii) intra-colony traveling waves of gene expression emerge across all genetic architectures tested, including constitutive reporter expression with no engineered regulatory network, and do so only after colonies stop growing; (iii) a nutrient-recovery model, in which depletion during growth is followed by diffusive replenishment after growth arrest, quantitatively recapitulates the observed wave dynamics; and (iv) a second inter-colony traveling wave propagates through the shared medium, consistent with a diffusible factor produced by the colonies suppressing gene expression at staggered times across the community. Our results support a model in which mechanical constraints organize colony growth and, combined with nutrient dynamics and subsequent changes in cellular metabolism after growth arrest, generate the observed traveling waves in gene expression.

## 2 Results

### 2.1 Colony expansion is edge-dominated

To characterize colony expansion and test the mechanical prediction of edge-dominated growth [19], we followed *E. coli* (MG1655) colonies on PDMS-agarose pads from single cells to millimeter-scale colonies by phase-contrast time-lapse microscopy every 10 min for up to 68 h (see Methods). Colonies were seeded using two inoculum concentrations, with the high-density condition tenfold more concentrated than the low-density condition (see Methods). Mechanical models predict that the local growth rate decays exponentially with distance from the colony edge [19]. Under radial symmetry, a corresponding radial velocity profile can be derived from this growth rate profile through biomass conservation (see supplementary section 1.1, Supplementary Fig. S1). We extracted colony displacement fields using mutual information image velocimetry and fitted the predicted radial velocity profile to the data (see supplementary section 1.2):

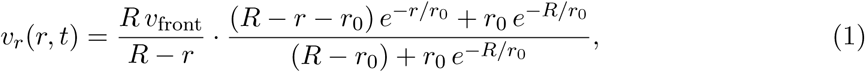

where *r* is the radial distance from the colony edge, *r*_0_ is the characteristic decay length of the radial growth-rate profile and provides a measure of the thickness of the actively growing region, *R*(*t*) is the colony radius, and *v*_front_ = *dR/dt* is the expansion speed of the front.

The model closely reproduced the observed velocity field (*R*^2^ = 0.892, Fig. 1A), showing that colony expansion is strongly edge-dominated. We then examined how the characteristic decay length *r*_0_ varied with colony size. Across all colonies, *r*_0_ increased with colony radius (Supplementary Fig. S2). Smaller colonies were excluded because the velocity profile model is ill-conditioned and fits poorly in the uniform expansion regime (see supplementary section 1.3, Supplementary Fig. S2). A weaker positive association persisted after restricting the analysis to colonies in the edge-dominated regime (*R >* 200 µm; Supplementary Fig. S3). Thus, within this regime, the absolute thickness of the actively growing region showed only a weak dependence on colony size. Consistent with this, *r*_0_*/R* decreased with colony radius (Fig. 1B), indicating that the actively growing region occupies a progressively smaller fraction of the colony as colony size increases. Because the model fits a single time-averaged *r*_0_ for each colony, this cross-colony analysis does not resolve how *r*_0_ changes during the growth of an individual colony.

**Figure 1:**
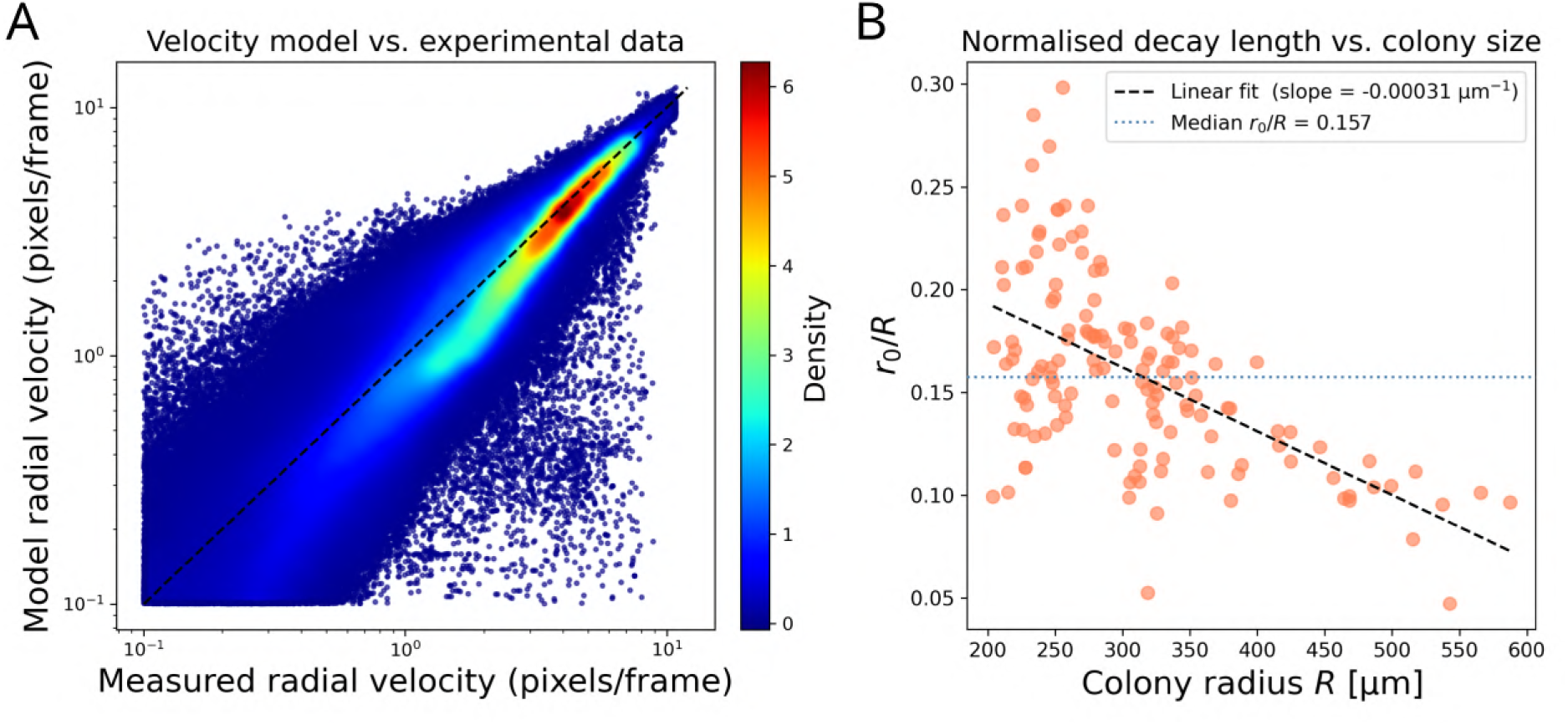
Edge-dominated colony expansion and its characteristic length scale. **(A)** Model radial velocity *v_r_* versus measured radial velocity from velocimetry, pooled across *n* = 152 colonies. Each point represents one grid window at one time point; color indicates local point density. The dashed line is the identity (1:1). Velocimetry measurements below *v_r_* = 0.1 pixels/frame were excluded because they fell below the reliable detection threshold (*R*^2^ = 0.892). **(B)** Normalized decay length *r*_0_*/R* as a function of colony radius *R*, for colonies in the edge-dominated growth regime (*R >* 200 µm, *n* = 139). The dashed line shows the decreasing trend (linear fit, slope = −3.1 × 10*^−^*^4^ µm*^−^*^1^); the dotted line indicates the median *r*_0_*/R* = 0.157. The decrease in *r*_0_*/R* indicates that the actively growing region occupies a smaller fraction of the colony as colony size increases within the edge-dominated regime. The direct relationship between *r*_0_ and *R* is shown in Supplementary Fig. S3.

### 2.2 Growth termination is density-dependent

Having characterized how growth is spatially organized within individual colonies, we next asked how colonies behave collectively, at the scale of the whole pad. Because every pad is identical in size, nutrient content, and preparation, the total resources available for growth are fixed by design. If colonies simply consume a shared, finite resource pool, the total biomass produced on a pad should then be the same regardless of how many colonies it contains.

We first characterized the growth of each colony by fitting its area expansion over time to a Gompertz model (see Methods), which provides the asymptotic final area *A_∞_*, the maximum growth rate *µ_m_*, the lag phase *λ*, and the growth stop time *t_s_*(Table 1). Colonies seeded at high density stopped growing earlier than those at low density, and across all pads the growth stop time decreased consistently as the number of colonies per pad increased (see supplementary section 1.4, Supplementary Fig. S4). Thus, growth termination was strongly shaped by colony density rather than determined solely by an intrinsic colony program.

**Table 1:**
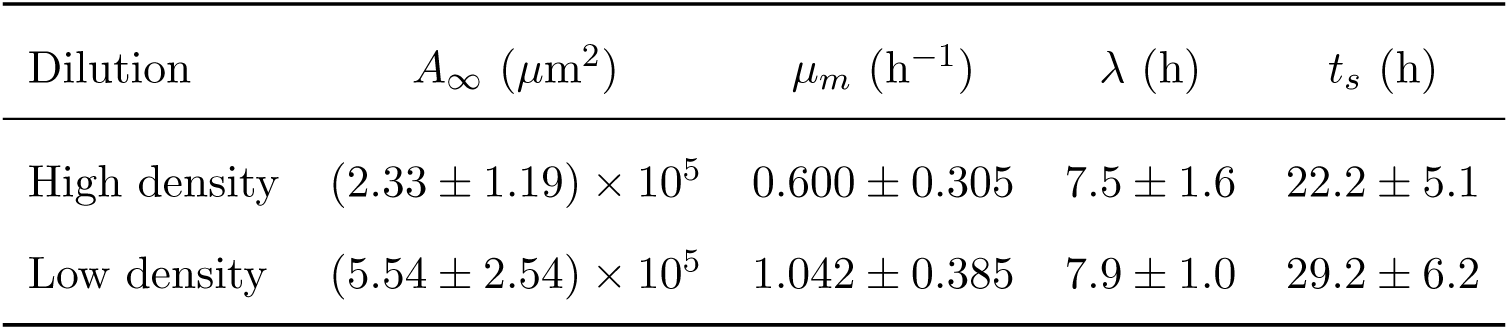
Gompertz growth model parameters for colonies at each seeding density. Values are mean ± SD (*n* = 160 colonies: 130 under high-density and 30 under low-density seeding conditions). All pairwise comparisons between dilutions were significant (Mann-Whitney *U*, *p <* 0.001). *A_∞_*: asymptotic final area; *µ_m_*: maximum growth rate; *λ*: lag phase; *t_s_*: growth stop time.

| Dilution | $A_\infty$ ($\mu\text{m}^2$) | $\mu_m$ ($\text{h}^{-1}$) | $\lambda$ (h) | $t_s$ (h) |
| --- | --- | --- | --- | --- |
| High density | $(2.33 \pm 1.19) \times 10^5$ | $0.600 \pm 0.305$ | $7.5 \pm 1.6$ | $22.2 \pm 5.1$ |
| Low density | $(5.54 \pm 2.54) \times 10^5$ | $1.042 \pm 0.385$ | $7.9 \pm 1.0$ | $29.2 \pm 6.2$ |

This crowding was also reflected in colony size. Individual colony area was larger at low seeding density than at high density (Fig. 2A). If colonies competed for a strictly fixed resource pool, mean colony area would be expected to decrease as 1*/N* (see supplementary section 1.6). Instead, it decreased more weakly with the number of colonies per pad, approximately as 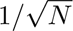 (Fig. 2B). Colonies are thus suppressed by their neighbors, but not as strongly as a simple division of a fixed total resource would imply.

**Figure 2:**
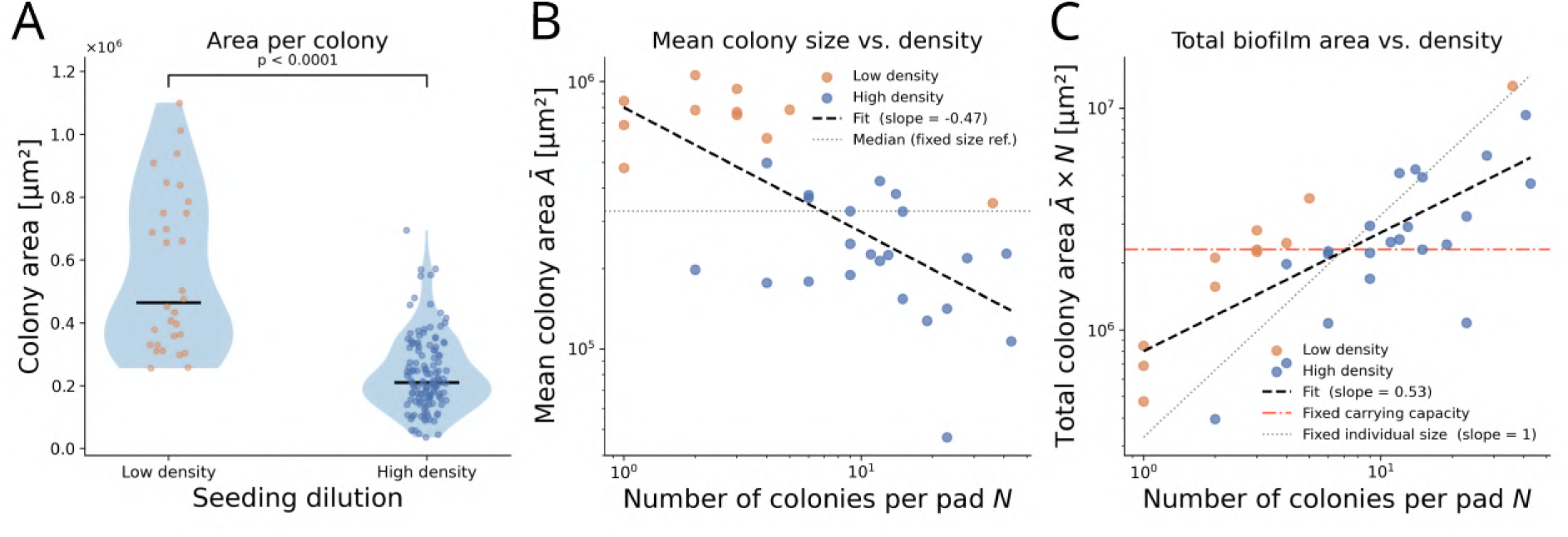
Density-dependent growth termination in a shared environment. **(A)** Final colony area under low- and high-density seeding conditions (*n* = 30 and *n* = 130 colonies, respectively). Violins show the distribution; points show individual colonies; horizontal lines indicate medians. Mann-Whitney *p <* 0.0001. **(B)** Mean colony area *Ā* per pad as a function of colony number *N*, shown in log-log space (*n* = 33 pads). The dashed line is a power-law fit (slope = −0.47); the dotted line indicates the median *Ā*. **(C)** Total colony area *Ā* × *N* per pad as a function of *N*, shown in log-log space. The dashed line is a power-law fit (slope = 0.53); the red dash-dot line is the fixed carrying capacity reference (*Ā* × *N* = const); the dotted line is the fixed individual size reference (slope = 1). Total colony area increases with *N* but more weakly than linearly, indicating partial competition for shared resources. Orange: low-density seeding; blue: high-density seeding; *n* = 33 pads.

The consequence of this scaling is most clearly seen at the pad level. If resources are fixed, total colony area should remain constant as the number of colonies increases. At first glance the data appear consistent with this, as total pad area was similar between the two seeding densities (see supplementary section 1.5, Supplementary Fig. S5A). However, when examined across the full range of colony numbers, total colony area did not remain flat but increased with colony number, scaling approximately as 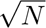 (Fig. 2C). This is inconsistent with colonies simply dividing a fixed resource pool, which would produce a constant total area, but total area also increased more weakly than the linear scaling expected if colonies grew independently without competition. The observed 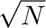 scaling lies between these two limits, indicating that colonies compete for shared resources but do not partition them exactly, consistent with competition acting locally rather than uniformly across the pad (see supplementary section 1.6).

Together, these results show that growth termination is not set by an intrinsic colony timer alone but is strongly shaped by the local environment that colonies share and collectively modify. As colonies grow, they deplete resources and alter their surroundings, and these density-dependent changes in the shared environment are closely tied to when growth stops. This raised the possibility that the same shared environment might shape not only growth, but also the spatial and temporal organization of gene expression.

### 2.3 A constitutive reporter reveals intra-colony traveling waves after growth arrest

Having established how growth is organized within colonies and how it responds to the collective environment, we next asked how gene expression evolves over the full course of colony development. We began with the simplest possible genetic system, a triple constitutive reporter [36] encoding three fluorescent proteins (mRFP1, EYFP, ECFP) under separate constitutive promoters with no engineered regulatory network (Fig. 3A, Supplementary Fig. S6). This circuit produces no oscillations and carries no regulatory architecture that could generate spatiotemporal patterns. Any spatiotemporal dynamics observed in this strain therefore cannot be produced by an engineered regulatory circuit.

**Figure 3:**
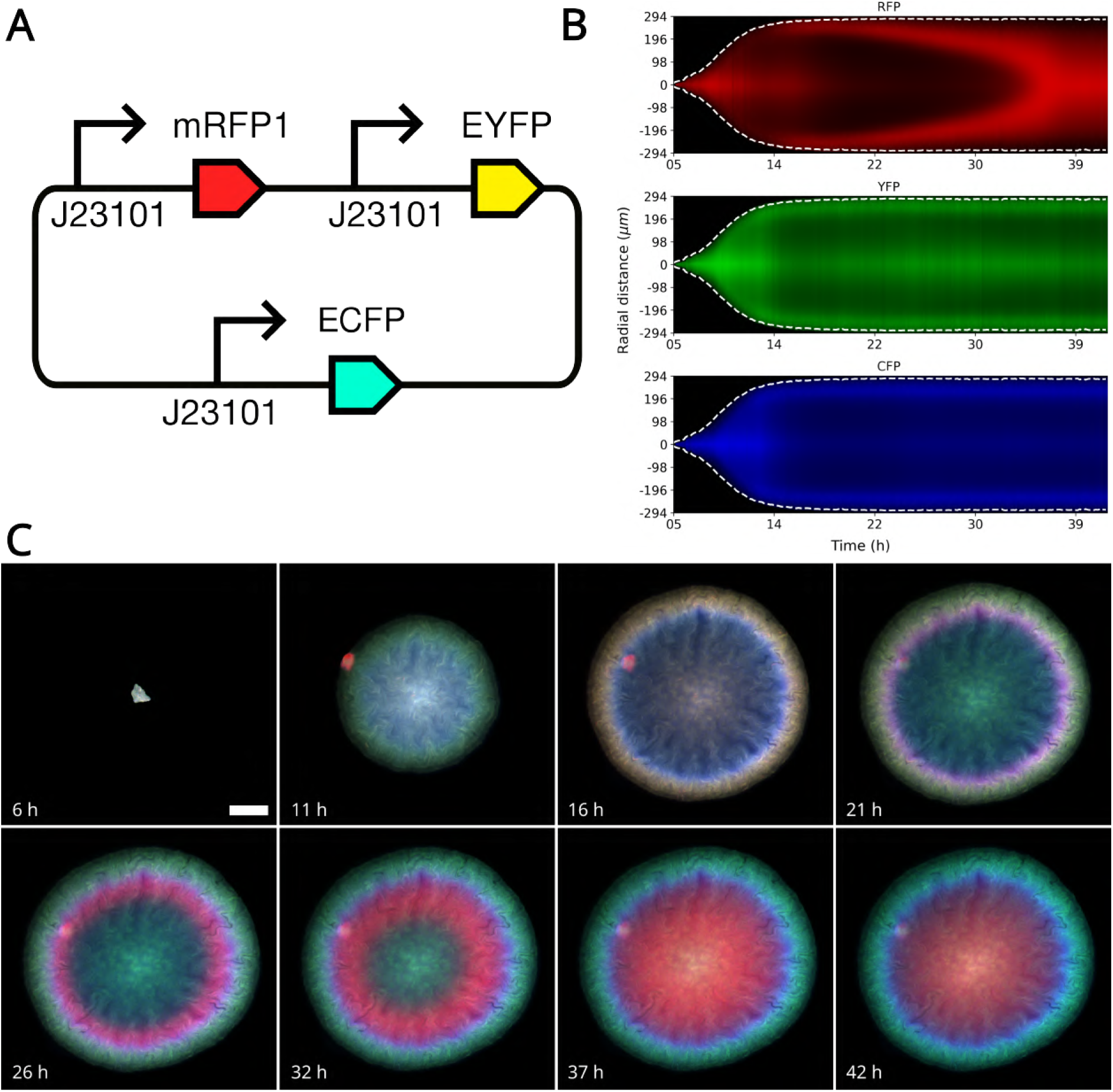
An intra-colony mRFP1 traveling wave emerges after growth arrest. **(A)** Schematic of triple constitutive reporter circuit: mRFP1 (red), EYFP (green), and ECFP (blue) are constitutively expressed under J23101 promoters with no engineered regulatory network. **(B)** Individual kymographs for RFP (top), YFP (middle), and CFP (bottom). The diagonal structure in RFP indicates a traveling wave; the patterns in YFP and CFP at the edge indicate static rings. Dashed white lines mark the colony boundary. **(C)** Time-lapse fluorescence images of a representative triple constitutive reporter colony at selected time points. The RFP wave initiates at the colony edge and propagates inward after growth arrest, while YFP and CFP remain at the periphery. Scale bar: 100 *µ*m.

Unexpectedly, we observed an intra-colony traveling wave of mRFP1 fluorescence that initiated at the colony edge and propagated inward toward the colony center, visible in time-lapse kymographs across multiple biological replicates and seeding conditions (Fig. 3B,C). EYFP and ECFP instead formed static rings at the colony edge with no inward propagation. The emergence of a traveling wave in a constitutive reporter shows that the wave does not require an engineered regulatory circuit and is instead consistent with a colony-level physiological or environmental origin.

The timing of the mRFP1 wave provides a key mechanistic clue, as it emerged at the colony edge only after the Gompertz growth stop time *t_s_*, was never observed during the active growth phase, and propagated inward over the subsequent 20–25 h (Fig. 3C). Together, these observations link wave initiation to the transition into growth arrest and point to a physiological or environmental change within the post-arrest colony as its origin. We therefore asked whether the same wave was also present in colonies carrying the repressilator, where circuit dynamics might modulate or interact with this environmental signal.

### 2.4 Intra-colony traveling waves generalize across circuit architectures

Having observed an intra-colony traveling wave in a circuit with no regulatory architecture, we asked whether the same post-arrest dynamics appeared in colonies carrying the repressilator and whether the circuit modulated or altered the wave. We tested two repressilator architectures carrying different reporter configurations.

In the single reporter repressilator [35], mVenus reports on the TetR node of the repressilator, while mCFP is constitutively expressed as an internal reference (Fig. 4A, Supplementary Fig. S7). Both channels showed traveling waves initiating at the colony edge and propagating inward after growth arrest (Fig. 4B,C). The constitutive mCFP is not regulated by the repressilator, yet it showed the same inward propagation as observed in the triple constitutive reporter, confirming that wave emergence does not require engineered regulatory dynamics. The regulated mVenus also showed a traveling wave after growth arrest, consistent with the environmental origin of the wave.

**Figure 4:**
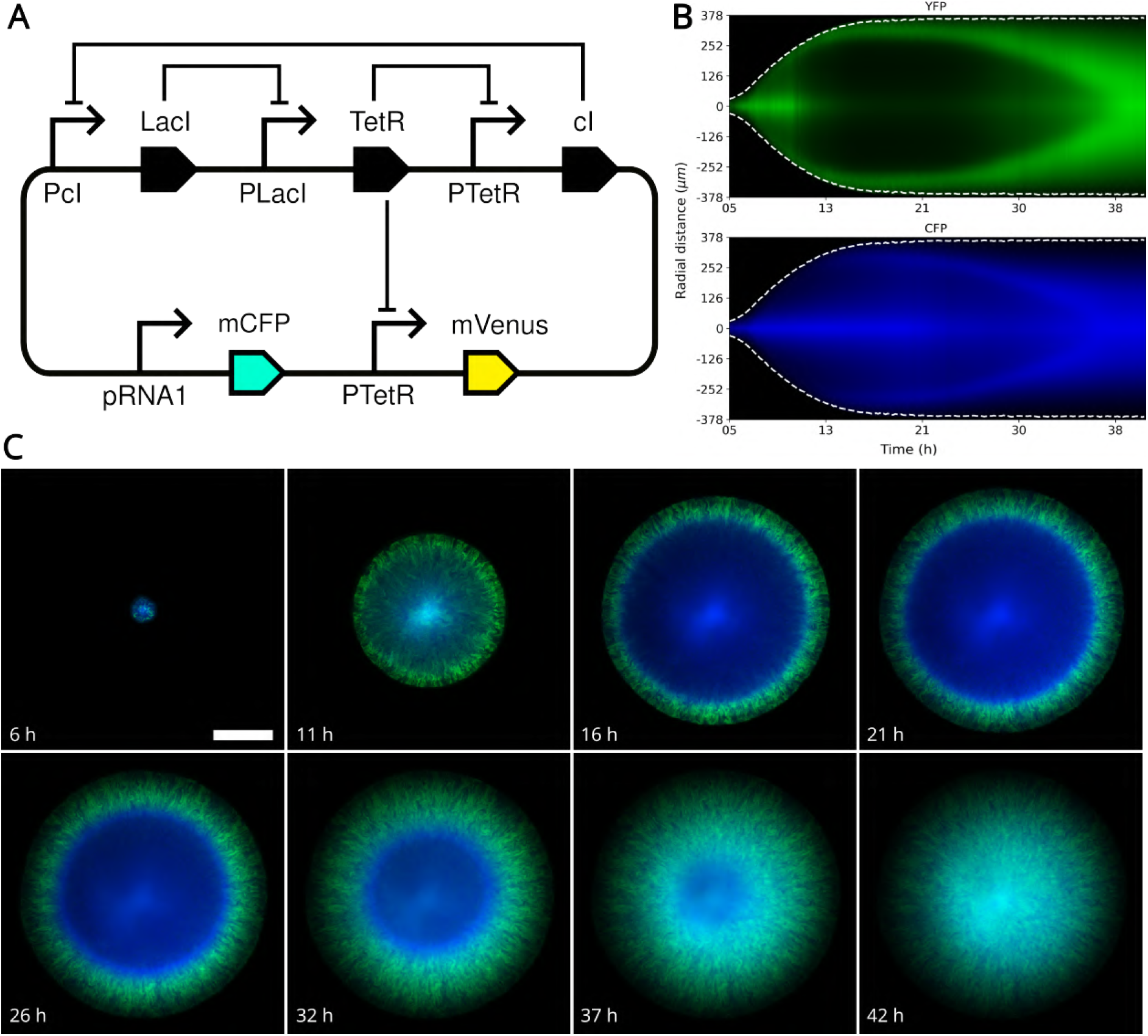
Intra-colony traveling waves emerge after growth arrest in the single reporter repressilator. **(A)** Schematic of the single reporter repressilator: mVenus (YFP, green) reports on the TetR node of the repressilator (LacI, TetR, cI), while mCFP (blue) is constitutively expressed as an internal reference. **(B)** Individual kymographs for mVenus (top) and mCFP (bottom). Both channels show a traveling wave after growth arrest. White dashed lines represent the edge of the colony. **(C)** Time-lapse fluorescence images of a representative single reporter repressilator colony at selected time points. Scale bar: 200 *µ*m.

In the triple reporter repressilator [35], all three nodes of the repressilator are reported: mKate2 (RFP) under *λ*cI, mVenus (YFP) under TetR, and mCFP under LacI (Fig. 5A, Sup-plementary Fig. S8). All three channels showed coordinated inward propagation after growth arrest, with phase differences between them consistent with their positions in the repressilator loop (Fig. 5B,C). Wave initiation was positively correlated with the Gompertz growth stop time *t_s_* across colonies, confirming that the wave onset is tied to the transition out of active growth rather than to a fixed experimental time.

**Figure 5:**
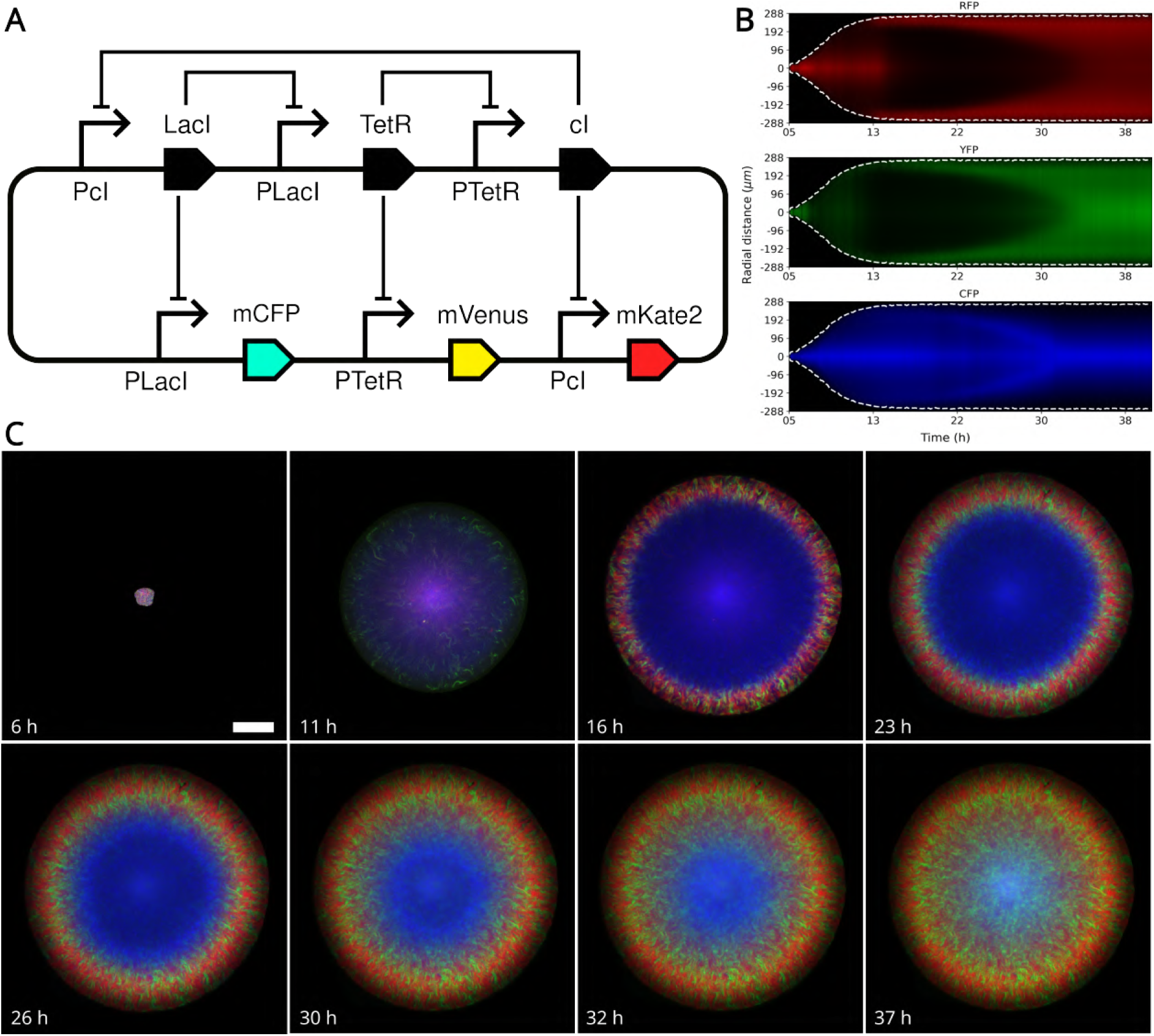
Intra-colony traveling waves emerge after growth arrest in the triple reporter repressilator. **(A)** Schematic of the triple reporter repressilator: mKate2 (RFP, red), mVenus (YFP, green), and mCFP (CFP, blue) are each regulated by their respective repressilator repressors (*λ*cI, TetR, and LacI). **(B)** Individual kymographs for mKate2 (top), mVenus (middle), and mCFP (bottom). All three reporters show the same inward wave after growth arrest, with phase differences between them consistent with their positions in the repressilator loop. **(C)** Time-lapse fluorescence images of a representative triple reporter repressilator colony at selected time points. Scale bar: 100 *µ*m.

Intra-colony waves appeared after growth arrest across constitutive and repressilator archi-tectures alike. This shared behavior suggests that a common colony-level process drives the dynamics independently of circuit topology. The wave appeared only after growth arrest and was never observed during active growth, linking its emergence to a post-arrest change in the colony environment. We therefore asked whether nutrient depletion during growth followed by diffusive recovery after arrest could account for the observed wave timing and propagation.

### 2.5 A nutrient-recovery model recapitulates intra-colony traveling waves after growth arrest

Given that waves after growth arrest were observed across both constitutive and repressilator architectures, we looked for a common physical explanation. The timing suggests a nutrient-driven process in which active growth depletes local nutrients faster than they can diffuse in from the surrounding agarose. When growth stops, nutrients from the surrounding medium diffuse back into the colony, generating a front of nutrient replenishment that propagates inward (Fig. 6A).

**Figure 6:**
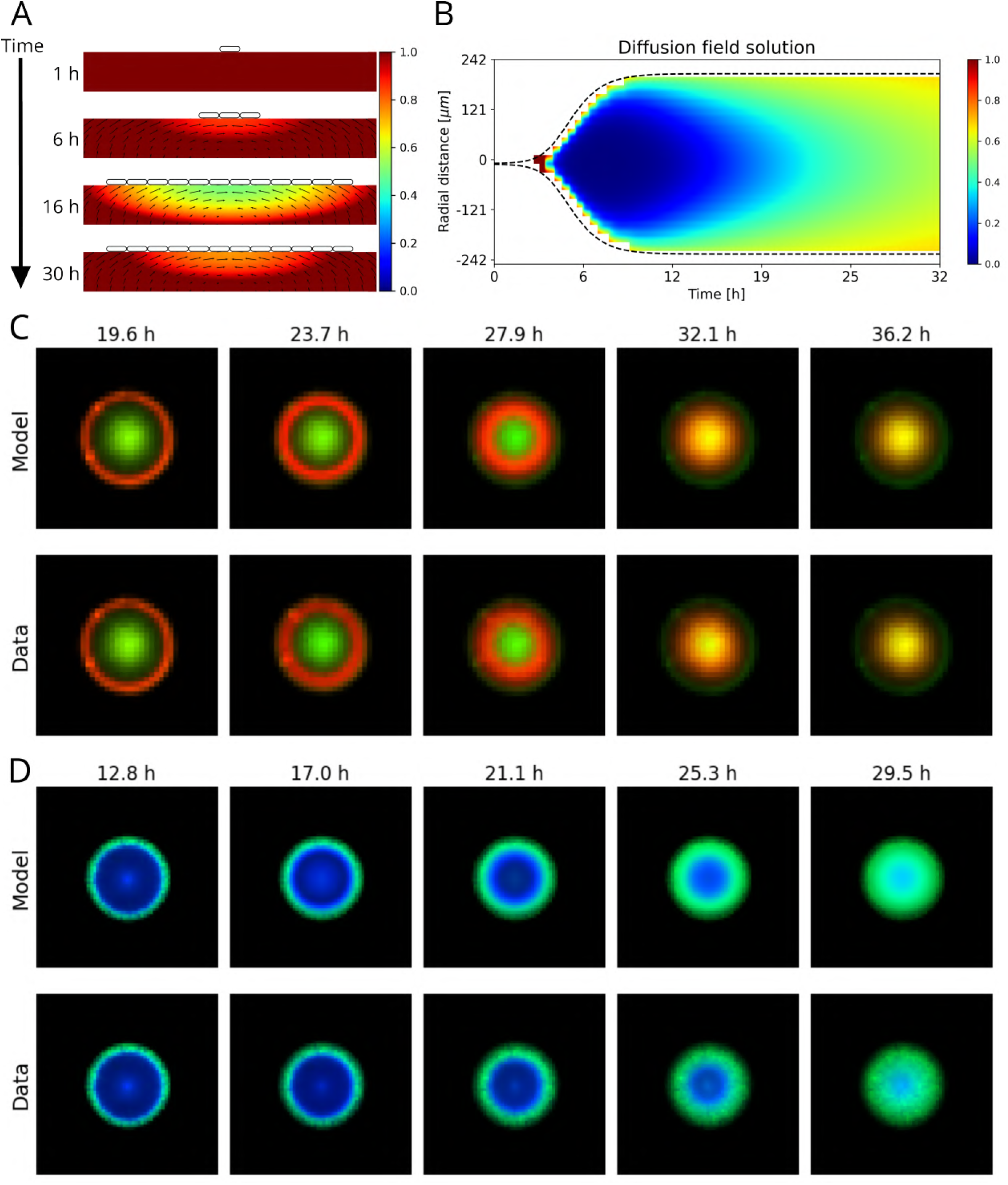
Nutrient-recovery model recapitulates traveling waves after growth arrest. **(A)** Conceptual model of nutrient depletion during growth and inward replenishment after growth arrest. **(B)** Simulated nutrient concentration field for a representative colony, showing depletion during active growth followed by recovery from the colony edge inward. Dashed lines mark the colony edge. **(C)** Model predictions (top) and experimental data (bottom) for a representative triple constitutive reporter colony at selected time points, showing the model reproduces the measured wave in the RFP channel. **(D)** Model predictions (top) and experimental data (bottom) for a representative single reporter repressilator colony, showing the regulated mVenus and constitutive mCFP channels, with the model reproducing the wave in both. Model simulations in **(C)** and **(D)** were initialized with the experimental fluorescence field at *t* = *t*_0_.

To formalize this, we constructed a model for nutrient dynamics within the colony domain. The nutrient concentration *S*(**x***, t*) evolves according to

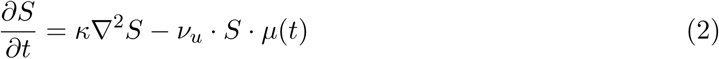

where *κ* is the coefficient of the diffusion term, interpreted here as an effective nutrient-propagation parameter, *ν_u_*is the nutrient uptake coefficient, and *µ*(*t*) is the time-dependent colony-level growth activity inferred from the colony expansion curve. In this model, nutrient recovery is driven by diffusion from the colony boundary, where the surrounding medium provides a higher nutrient concentration after growth arrest.

If gene expression is coupled to local nutrient availability, this front would produce a traveling wave of increased reporter expression moving from the colony edge toward the center. Thus, we represent reporter fluorescence *F_i_*(**x***, t*) as depending on the nutrient concentration

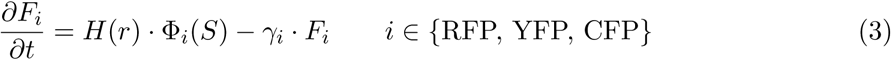

where *γ_i_*is the reporter turnover rate, Φ*_i_*(*S*) is the nutrient-dependent production rate, and *H*(*r*) represents the radial height profile.

Gene expression is coupled to the local nutrient field through a production rate represented by a nutrient-dependent Hill function

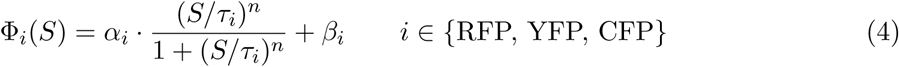

where *α_i_* is the nutrient-dependent production amplitude, *τ_i_* is the half-saturation constant of the nutrient-dependent response, *n* is the Hill coefficient, and *β_i_* is the basal expression level. Therefore, the maximum total production rate is *α_i_* + *β_i_*.

Although the colonies grow mostly in two dimensions, they are not quite flat. Cells can accumulate along the z-axis, especially near the colony center, increasing the measured fluorescence intensity in those regions. To account for this, we included a fixed radial weighting term in the fluorescence model representing this height profile *H*(*r*)

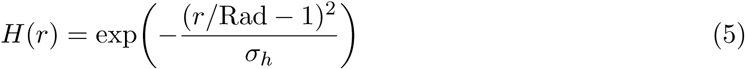

where *r* is the distance from the colony edge towards the center, Rad is the maximum value of this distance within the colony (equivalent to the colony radius), and *σ_h_*is a dimensionless parameter that controls the radial spread of the height profile in normalized colony coordinates. Thus, *H*(*r*) is highest near the colony center and smaller near the edge.

For the single reporter repressilator, the same nutrient field *S*(**x***, t*) and height profile *H*(*r*) were coupled to the repressilator architecture. The topology follows a three-node inhibitory ring,

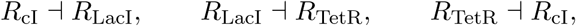

where TetR also represses the fluorescent reporter mVenus,

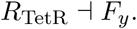

The repressor dynamics are given by

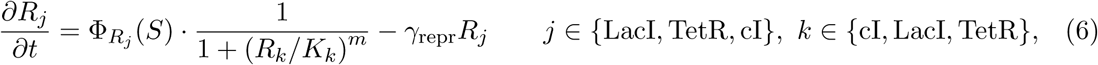

where *k* is the repressor of *j* following the same ordering, *K_k_* is the repression half-saturation constant, *γ*_repr_ is the repressor turnover rate, and *m* is the repression Hill exponent. Since the mVenus reporter is repressed by TetR, the dynamics of mVenus are

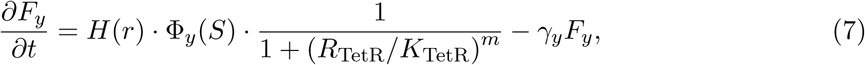

where *γ_y_*is the mVenus turnover rate, whereas the CFP reporter is unrepressed and its dynamics are

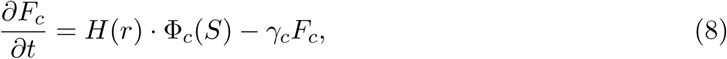

with *γ_c_* the CFP turnover rate. The nutrient-dependent production terms for the repressors are

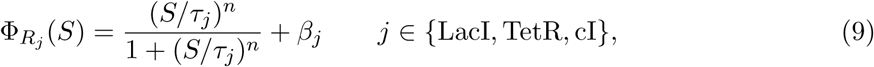

where *τ_j_* is the nutrient half-saturation constant and *β_j_* is the basal production level of repressor *j*. For the mVenus reporter, the nutrient response uses the *R*_cI_ nutrient-response parameters,

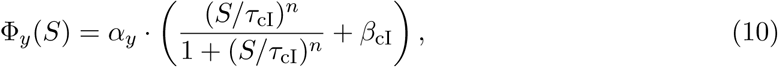

where *α_y_* is the nutrient-dependent production amplitude. For the CFP reporter,

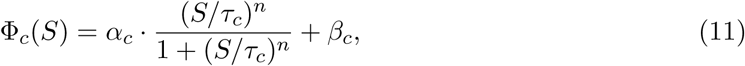

where *α_c_*is the nutrient-dependent production amplitude, *τ_c_* is the nutrient half-saturation constant, and *β_c_*is the basal production level.

We implemented the model on a 2D grid for each colony individually, using an idealized circular geometry derived from the colony expansion curve, and fitted it to the experimental two-dimensional fluorescence image stacks by nonlinear least-squares optimization (see Methods). We fitted the model separately to seven triple constitutive reporter colonies and eight single reporter repressilator colonies and present one representative fit for each architecture in Fig. 6. Solving the nutrient dynamics produces a field that recovers from the colony edge inward after growth arrest, shown for the representative triple constitutive reporter colony in Fig. 6B.

For the triple constitutive reporter, the model reproduced the measured wave in the RFP channel (Fig. 6C). For the single reporter repressilator, the nutrient field and height profile were coupled to the repressilator ring, with TetR repressing the mVenus reporter, and the model reproduced the wave in both the regulated mVenus and the constitutive mCFP channels (Fig. 6D). Fitted parameters for the representative colonies are listed in Table 2 and Table 3, and per-colony fit residuals are reported in Supplementary Table S2.

**Table 2:** Parameters of the triple constitutive reporter nutrient-recovery model fitted to the representative colony.

| Parameter | Description | Units | Value |
| --- | --- | --- | --- |
| $\kappa$ | Effective nutrient propagation coefficient | $\text{m}^2 \text{s}^{-1}$ | $1.33 \times 10^{-13}$ |
| $\nu_u$ | Nutrient uptake coefficient | dimensionless | 9.35 |
| $\gamma_r$ | RFP turnover rate | $\text{s}^{-1}$ | $3.15 \times 10^{-4}$ |
| $\gamma_y$ | YFP turnover rate | $\text{s}^{-1}$ | $7.34 \times 10^{-6}$ |
| $\alpha_r$ | RFP nutrient-dependent production amplitude | a.u. $\text{s}^{-1}$ | 9.91 |
| $\alpha_y$ | YFP nutrient-dependent production amplitude | a.u. $\text{s}^{-1}$ | $7.74 \times 10^{-6}$ |
| $\beta_r$ | RFP basal expression level | a.u. $\text{s}^{-1}$ | 0.534 |
| $\beta_y$ | YFP basal expression level | a.u. $\text{s}^{-1}$ | $1.70 \times 10^{-3}$ |
| $\tau_r$ | RFP nutrient half-saturation constant | dimensionless | 0.430 |
| $\tau_y$ | YFP nutrient half-saturation constant | dimensionless | 2.52 |
| $n$ | Nutrient-response Hill coefficient | dimensionless | 22.2 |
| $\sigma_h$ | Width of radial height profile | dimensionless | 0.300 |
| RFP half-life | Derived from $\gamma_r$ | h | 0.61 |
| YFP half-life | Derived from $\gamma_y$ | h | 26.2 |

**Table 3:** Parameters of the single reporter repressilator nutrient-recovery model fitted to the representative colony. One colony was excluded from analysis due to biologically implausible K_LacI_ value and highest MSE among fitted colonies.

| Parameter | Description | Units | Value |
| --- | --- | --- | --- |
| $\kappa$ | Effective nutrient propagation coefficient | $\text{m}^2 \text{s}^{-1}$ | $2.07 \times 10^{-13}$ |
| $\nu_u$ | Nutrient uptake coefficient | dimensionless | 1.27 |
| $\sigma_h$ | Width of radial height profile | dimensionless | 0.602 |
| $K_{\text{LacI}}$ | LacI repression half-saturation constant | a.u. | 0.0759 |
| $K_{\text{TetR}}$ | TetR repression half-saturation constant | a.u. | 50.5 |
| $K_{\text{cI}}$ | cI repression half-saturation constant | a.u. | 2.89 |
| $\gamma_{\text{repr}}$ | Repressor turnover rate | $\text{s}^{-1}$ | $3.87 \times 10^{-5}$ |
| $\gamma_y$ | mVenus turnover rate | $\text{s}^{-1}$ | $6.36 \times 10^{-6}$ |
| $\gamma_c$ | CFP turnover rate | $\text{s}^{-1}$ | $2.55 \times 10^{-4}$ |
| $\alpha_y$ | mVenus nutrient-dependent production amplitude | a.u. $\text{s}^{-1}$ | 0.561 |
| $\alpha_c$ | CFP nutrient-dependent production amplitude | a.u. $\text{s}^{-1}$ | 6.20 |
| $\beta_{\text{LacI}}$ | LacI basal production level | a.u. $\text{s}^{-1}$ | $2.35 \times 10^{-6}$ |
| $\beta_{\text{TetR}}$ | TetR basal production level | a.u. $\text{s}^{-1}$ | $2.59 \times 10^{-3}$ |
| $\beta_{\text{cI}}$ | cI basal production level | a.u. $\text{s}^{-1}$ | $2.55 \times 10^{-2}$ |
| $\beta_c$ | CFP basal expression level | a.u. $\text{s}^{-1}$ | 2.05 |
| $\tau_{\text{LacI}}$ | LacI nutrient half-saturation constant | dimensionless | 0.817 |
| $\tau_{\text{TetR}}$ | TetR nutrient half-saturation constant | dimensionless | 0.766 |
| $\tau_{\text{cI}}$ | cI nutrient half-saturation constant | dimensionless | 0.519 |
| $\tau_c$ | CFP nutrient half-saturation constant | dimensionless | 0.474 |
| $n$ | Nutrient-response Hill coefficient | dimensionless | 25.4 |
| $rep0_{\text{LacI}}$ | Initial LacI concentration | a.u. | 0.0845 |
| $rep0_{\text{TetR}}$ | Initial TetR concentration | a.u. | $2.43 \times 10^{-3}$ |
| $rep0_{\text{cI}}$ | Initial cI concentration | a.u. | $1.69 \times 10^{-3}$ |
| Repressor half-life | Derived from $\gamma_{\text{repr}}$ | h | 4.97 |
| mVenus half-life | Derived from $\gamma_y$ | h | 30.3 |
| CFP half-life | Derived from $\gamma_c$ | h | 0.756 |

In the triple constitutive reporter only the RFP channel travels as a wave, while YFP and CFP stay close to flat. The fitted RFP amplitude supports this, since the nutrient-dependent amplitude of RFP (*α_r_* = 9.9) is large relative to its basal level (*β_r_*= 0.53), so RFP production is strongly modulated by the nutrient front. The static YFP channel was reproduced with a negligible nutrient-dependent amplitude (*α_y_*≈ 8 × 10*^−^*^6^), although this value was held at its initial value and not independently informed by the fit (Table S3). CFP was assumed to share the same production parameters as YFP and was not fitted separately.

The repression Hill exponent was fixed at *m* = 4 for all fits. In the single reporter repressilator both fluorescent channels travel, and the fitted amplitudes support this, since both mVenus (*α_y_* = 0.56) and mCFP (*α_c_* = 6.2) carry a nutrient-dependent contribution comparable to or larger than their basal production, so both reporters follow the nutrient front. The fitted turnover rates differ substantially between channels. The effective turnover was fast for RFP and CFP, with values below one hour, and slow for YFP and mVenus, with values above 24 hours. We interpret these as effective fluorescence turnover rates rather than measured protein half-lives, because the fitted *γ* can absorb fluorophore maturation, photobleaching and other reporter-specific effects in addition to protein degradation. The difference between fast and slow channels nonetheless shapes whether a reporter tracks the nutrient front or accumulates as a static pattern. In both models the fitted nutrient-response Hill coefficient was high (*n* ≈ 22 for the triple constitutive reporter and *n* ≈ 25 for the single reporter repressilator), indicating that the nutrient response acts as a sharp switch rather than a graded function.

The fitted values of *κ* (1.3 × 10*^−^*^13^ m^2^ s*^−^*^1^ for the triple constitutive reporter and 2.1 × 10*^−^*^13^ m^2^ s*^−^*^1^ for the single reporter repressilator) are several orders of magnitude below molecular diffusion coefficients for small nutrients in agarose-like media [12, 37], so *κ* acts as an effective propagation parameter rather than a physical diffusivity. With this interpretation the model captures the timing of wave initiation at the colony edge and its inward propagation.

These results support nutrient depletion during growth followed by diffusive replenishment after arrest as a common mechanism for the waves observed across circuit architectures. Colonies on a shared pad are not isolated, since they grow in a common medium through which signals can diffuse between colonies. We therefore next asked whether the same diffusive coupling could produce a second wave spanning multiple colonies at the pad scale.

### 2.6 An inter-colony suppression wave suggests diffusive coupling

Having identified an intra-colony wave after growth arrest, we next asked whether reporter expression dynamics were also coordinated between colonies sharing the same agarose pad. At approximately 50 h after seeding, we observed a second, qualitatively distinct inter-colony suppression wave, in which reporter fluorescence decreased across the pad and the suppression reached different colonies at different times (Fig. 7A). Unlike the nutrient-recovery wave, which propagated inward within each colony, this wave traveled directionally across the pad and was associated with a decrease rather than an increase in reporter signal.

**Figure 7:**
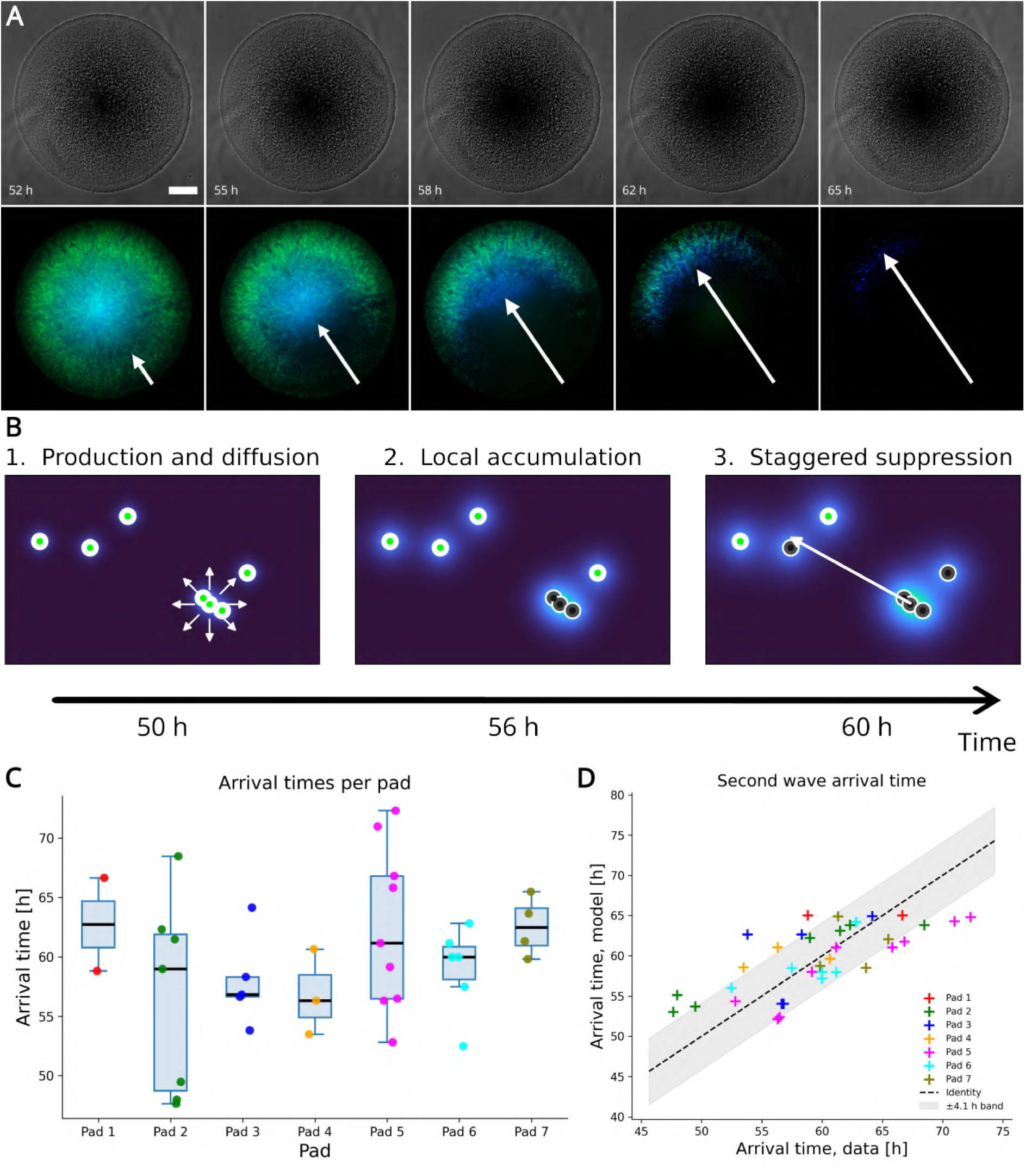
Inter-colony suppression dynamics are consistent with diffusive coupling. **(A)** Representative late-time phase-contrast (top) and fluorescence (bottom) images of a single reporter repressilator colony. Arrows indicate the direction of asymmetric fluorescence suppression across the colony, consistent with a signal approaching through the shared medium. Scale bar: 100 *µ*m. **(B)** Inter-colony model schematic. Colonies continuously produce a diffusible factor that spreads through the shared medium. Panels 1–3 depict production and diffusion, local accumulation and early suppression, and staggered threshold crossing; the arrow indicates suppression progression. Green-centered and dark-centered markers indicate colonies below and above the suppression threshold, respectively. **(C)** Inter-colony suppression-wave arrival times across seven pads. Boxes show medians and interquartile ranges; colored points show individual colonies. **(D)** Simulated versus observed arrival times. The dashed line is the identity and the shaded region indicates ±4.1 h RMSE. Pearson *r* = 0.703, *p* = 1.7 × 10*^−^*^6^, *n* = 36.

These observations suggested a mechanism in which colonies continuously produce a diffusible factor that accumulates and spreads through the shared medium and crosses a suppression threshold at different positions over time (Fig. 7B). Within each pad, colonies showed staggered suppression times rather than a simultaneous decrease, with the fluorescence decrease arriving at different colonies depending on their position within the pad. Across replicate pads from the same experiment, the phenomenon occurred within a similar time window, suggesting that it reflected a reproducible developmental process rather than an experimental artifact (Fig. 7C, Supplementary Fig. S9).

To test whether diffusive propagation could account for this timing, we used a minimal model in which each colony was represented as a point source of an unidentified diffusible factor *q*(**x***, t*), whose concentration evolves according to

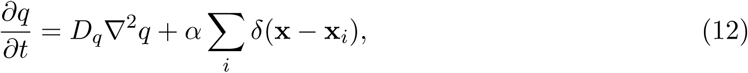

where *D_q_*is the effective diffusion coefficient, *α* is the common source strength, and **x***_i_* is the position of colony *i*. Treating colonies as point sources is a minimal approximation. In the numerical implementation, each colony was represented as a localized source on the simulation grid (see Methods). The predicted arrival time at each colony position was defined as the time at which the local concentration first exceeded a threshold *q*_thresh_.

The source strength was fixed to *α* = 1 for all colonies and decay of the factor was neglected. Because the model was used to test relative differences in arrival time between colony positions rather than the absolute onset of factor accumulation, a single global offset was estimated as the mean difference between observed and simulated times and added to all simulated arrival times. Since the diffusion equation is linear in the source strength, *α* and *q*_thresh_ are not separately identifiable, so *q*_thresh_ is reported in simulation units. The fitted parameters are listed in Table 4. Model predictions for arrival times were consistent with the experimentally observed suppression times (Fig. 7D), supporting an interpretation based on diffusion. The identity of the diffusible factor remains to be determined, but the spatial dependence of arrival times across the pad is consistent with a metabolic byproduct or other soluble signal diffusing through the shared medium.

**Table 4:** Parameters of the inter-colony diffusion model fitted jointly across the seven pads (*n* = 36 colonies). *D_q_* and *q*_thresh_ were the only estimated parameters. The common source strength was fixed to *α* = 1 and was identical for all colonies. Because the diffusion equation is linear in the source strength, the threshold and source strength are not separately identifiable, so *q*_thresh_ is reported in simulation units. The fit tested relative differences in arrival time between colony positions, and a single global offset *t*_offset_ was estimated as the mean of observed minus simulated times and added to the simulated arrival times for absolute alignment. Model and observed arrival times agreed with a root mean square error of 4.1 h (Pearson *r* = 0.703, *p* = 1.7 × 10*^−^*^6^, *n* = 36).

| Parameter | Description | Units | Value |
| --- | --- | --- | --- |
| $D_q$ | Effective diffusion coefficient | $\text{m}^2 \text{s}^{-1}$ | $7.32 \times 10^{-12}$ |
| $\alpha$ | Common source strength, fixed and identical across colonies | sim. units | 1 |
| $q_{\text{thresh}}$ | Suppression threshold | sim. units | $6.91 \times 10^{10}$ |
| $t_{\text{offset}}$ | Global temporal offset added to simulated arrival times | h | 49.5 |

Together, the intra-colony nutrient-recovery wave and the inter-colony suppression wave suggest that reporter expression dynamics in developing colonies are shaped by diffusive processes at multiple spatial scales, from local nutrient dynamics within individual colonies to coupling between colonies through the shared medium.

## 3 Discussion

In bacterial colonies carrying engineered genetic constructs, gene expression is often interpreted as a direct output of the circuit, yet colony development continuously reshapes the environment in which that circuit operates [33]. By following founding colonies from single cells to millimeter-scale colonies, we identified a developmental sequence in which growth modifies the mechanical and chemical environment and these self-generated changes later feed back on gene expression [11]. Mechanics organized colony expansion, colony density shaped the timing of growth arrest, the intra-colony wave was consistent with post-arrest environmental recovery, and the later inter-colony suppression wave suggested diffusive coupling through the shared medium. Together, these results support a model in which colonies generate spatial information through growth, metabolism, and diffusion rather than simply reading externally imposed gradients or circuit-encoded spatial programs [21]. This developmental sequence illustrates how nonlinear feedback between growth, environmental change, and gene expression can organize traveling waves across spatial scales [8].

Colony expansion followed the edge-dominated pattern predicted from mechanical constraints, and the measured radial velocity field was well described by the corresponding profile [13, 16–19, 38]. However, the decrease in *r*_0_*/R* with colony radius shows that a single average decay length per colony is a simplification, and models in which *r*_0_ varies with size, density, or local stress may better capture the transition from expansion to growth arrest [37, 39, 40]. Growth termination was also inconsistent with a fixed pad-level carrying capacity. The complementary scaling of individual and total colony area with colony number was consistent with partial local competition rather than strict partitioning of a fixed global resource pool, indicating that growth arrest is density-dependent and shaped by a shared but spatially structured microenvironment. Founder density and spatial configuration are known to alter pattern formation and competitive outcomes in colony biofilms, while resource limitation can itself drive spatial organization [41–44]. In this sense, arrest is not merely the end of colony expansion but a transition into a new environmental regime.

This transition was most clearly revealed by the constitutive reporter, in which an inward mRFP1 wave emerged only after growth had stopped and was never observed during active expansion. Because this construct contains no engineered regulatory interaction expected to encode such a pattern, the wave points to a physiological or environmental change associated with the post-arrest colony state. A minimal nutrient-recovery model recapitulated the timing and inward propagation observed in the constitutive reporter and single reporter repressilator by assuming that, during expansion, consumption depletes a limiting variable within the colony faster than diffusion can replenish it [21, 37]. After growth arrest, reduced consumption allows recovery from the edge inward, while reporter expression coupled to local metabolic state converts this recovery front into a traveling wave. The fitted parameter *κ* should be interpreted as a phenomenological effective propagation parameter rather than a literal molecular diffusion coefficient because it combines transport, uptake, and delayed fluorescence responses [12, 37]. The high fitted Hill coefficient further suggests a nonlinear, switch-like coupling that converts gradual environmental recovery into a sharp change in reporter expression once local conditions cross an effective threshold. This interpretation is consistent with the broader role of diffusion and nutrient gradients in structuring biofilm physiology [12, 15, 20, 45], although the molecular identity of the relevant nutrient or physicochemical variable remains unresolved.

Across the repressilator architectures, the common post-arrest dynamics were filtered differently by circuit topology and reporter properties. In the single reporter repressilator, both regulated mVenus and constitutive mCFP displayed inward waves, whereas in the triple reporter repressilator all three reporters propagated with temporal offsets consistent with the regulatory architecture. These observations separate a common colony-level driver from the circuit- and reporter-specific form of its readout. In the constitutive construct, mRFP1 traveled while EYFP and ECFP formed comparatively static, edge-enriched patterns. These differences may arise from reporter-specific maturation [46], effective accumulation and turnover [47], or sensitivity of expression to cellular physiology [33]. Fluorescent reporters therefore do not measure transcriptional activity instantaneously but temporally filter the underlying dynamics through production, maturation, degradation, and dilution [47]. The observed patterns consequently reflect the combined effects of circuit architecture, reporter behavior, cellular physiology, and the changing colony environment.

At later stages, an inter-colony suppression wave propagated asymmetrically across the pad and reached different colonies at staggered times. A minimal model in which colonies acted as sources of an unidentified diffusible factor reproduced the spatial dependence of these arrival times, suggesting coupling through the shared medium even without an engineered communication system [25]. Metabolic stress can also be communicated through propagating potassium waves within bacterial biofilms, and these electrical signals can influence cells outside the producing community [9, 48, 49]. The responsible factor could be a metabolic byproduct, a pH-related chemical change, or another soluble molecule produced during colony maturation. As with *κ*, the fitted *D_q_* should be regarded as an effective propagation parameter rather than a measured molecular diffusion coefficient. Together, these observations extend environmental feedback from processes organizing gene expression within individual colonies to chemical coupling across the wider community.

Many synthetic patterning systems generate spatial organization through engineered oscillators [35, 50], bistable switches [51, 52], Turing circuits [53, 54], optogenetic forcing [50], or programmed intercellular communication [55–59]. These include concentric rings formed by the improved repressilator [35], spatial patterns produced by optogenetic forcing [50], and growth-dependent fate bifurcations arising from interactions between colony-generated nutrient gradients and regulatory architecture [52]. Natural biofilms can also translate temporal gene expression dynamics into spatial differentiation through stochastic pulsing and clock and wavefront mechanisms [10, 60]. Here, by contrast, the intra-colony wave emerged after growth arrest and in a constitutive reporter, consistent with post-arrest environmental recovery rather than requiring an engineered regulatory instability. Because similar endpoint patterns can arise through different developmental mechanisms [11], time-resolved measurements are essential for distinguishing patterns formed during active expansion from those generated after growth stops.

The models used here are intentionally minimal and therefore define the scope of the mechanistic conclusions. The nutrient-recovery model represents an effective limiting variable rather than a specific nutrient and does not explicitly resolve oxygen, pH, waste accumulation, or three-dimensional transport. Its threshold-like response is phenomenological, the triple reporter repressilator was not modeled quantitatively, and the shared environment model treats colonies as sources of an unidentified diffusible field. These simplifications motivate perturbing medium composition, transport conditions, colony spacing, and reporter identity to distinguish alternative environmental mechanisms and identify the molecular species involved. More broadly, the environment should be treated as a design variable in synthetic biofilm engineering rather than as background noise around a genetic circuit [11]. This becomes increasingly important as systems are scaled from isolated microcolonies to multi-colony environments, patterned consortia, or engineered living materials [61], where local feedback and inter-colony coupling can reshape circuit output [55, 59]. Related processes may also contribute to physiological heterogeneity, metabolic microniches, and stress-tolerant subpopulations in natural biofilms [15, 20, 62–64]. Together, our results show that engineered gene circuits in bacterial colonies operate within a self-modifying environment that can generate spatial information across colony and community scales. The colony environment is therefore not merely a background condition or boundary constraint but an active component of the pattern-generating system.

## 4 Methods

### 4.1 Bacterial strains and plasmids

*E. coli* strains MG1655 and MC4100 were used in this study. Three plasmids were used:

- **Triple constitutive reporter (pAAA):** encoding mRFP1, EYFP, and ECFP under separate J23101 constitutive promoters (p15A origin) [36]. Used with MG1655.
- **Single reporter repressilator (pLPT20)** (Addgene #85460): encoding LacI, TetR, and *λ*cI in a feedback loop; mVenus (YFP) regulated by pTet and mCFP constitutively expressed from pRNA1 (pSC101 origin) [35]. Used with MC4100.
- **Triple reporter repressilator (pLPT107)** (Addgene #85525): with mKate2 regulated by p*λ*cI, mVenus regulated by pTet, and mCFP regulated by pLac (pSC101 origin [35]). Used with MC4100.

Overnight cultures were grown in LB at 37*^◦^*C with shaking at 250 rpm for 12–16 hours, supplemented with kanamycin (50 *µ*g ml*^−^*^1^) and/or carbenicillin (100 *µ*g ml*^−^*^1^) as appropriate. Cultures were diluted 10*^−^*^5^- or 10*^−^*^6^-fold in LB for seeding onto agarose pads, corresponding to the high- and low-density seeding conditions, respectively. The repressilator strains were grown in M9-CasAA supplemented with 10% LB.

### 4.2 PDMS-agarose imaging system

Agarose pads (1.5% w/v low-melt agarose in M9 minimal medium supplemented with LB for repressilator strains) were cast within PDMS frames (external 15 × 20 × 3 mm) placed on 1-mm glass slides. Three microliters of diluted culture was seeded on each pad. Pads were sealed with a #1.5 coverslip. The PDMS frame–coverslip assembly limited sample dehydration, while the gas permeability of PDMS supported oxygen exchange, enabling stable aerobic growth for up to 72 hours at 37*^◦^*C without condensation or dehydration artifacts. Up to 40 positions per pad were imaged simultaneously.

### 4.3 Microscopy

Time-lapse fluorescence and phase-contrast microscopy was performed on a Nikon Eclipse TiE inverted microscope with automated XY stage (Nikon Ti-S-ER) and Nikon Perfect Focus System. Images were acquired using a Photometrics Prime B sCMOS camera. Objectives: 10× (NA 0.30), 20× (NA 0.75), 40× (NA 0.60), 60× (NA 1.4 oil), 100× (NA 1.3 oil). Fluorescence channels: CFP (434 ± 8.5 nm excitation, 479 ± 20 nm emission), YFP (497 ± 8 nm excitation, 535 ± 11 nm emission), RFP/mCherry-A (578 ± 10.5 nm excitation, 641 ± 37.5 nm emission). Images were captured every 10 minutes. Temperature was maintained at 37*^◦^*C using an Okolab air heater. All images were acquired as 16-bit grayscale.

### 4.4 Image analysis

All image processing was performed in Python using NumPy [65], SciPy [66], Scikit-image [67], Pandas [68], and Matplotlib [69]. Colony boundaries were tracked using an iterative active contour (snake) algorithm [70] that propagates from the previous time step, minimizing an energy function balancing elasticity, bending stiffness, and gradient-based external forces. Only circular or near-circular colonies were kept for quantitative analysis. Velocity fields were computed using a mutual information-based particle image velocimetry algorithm [71–73] in which displacement vectors were estimated by maximizing the mutual information between small interrogation windows in consecutive frames. This approach is robust to the nonlinear intensity variations and overlapping structures characteristic of phase-contrast colony images. Kymographs were computed by dividing each colony mask into 16-pixel-wide radial bands from the edge inward and averaging the quantity of interest within each band at each time point. Wave starting and arrival times were identified as the time of the peak fluorescence value at the colony edge and colony center, respectively.

### 4.5 Gompertz growth model

Colony area *A*(*t*) was fitted to the Gompertz model [74]:

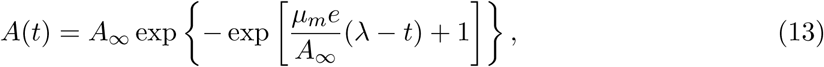

where *A_∞_* is the asymptotic maximum area, *µ_m_* is the maximum area expansion rate, and *λ* is the lag phase. The time of maximum growth rate was *t_m_* = *λ* + *A_∞_/*(*µ_m_e*), and growth stop time *t_s_* was defined as the time when *dA/dt <* 10 pixels (approximately 10 *µ*m^2^h*^−^*^1^). Fitting was performed using scipy.optimize.curve fit.

### 4.6 Nutrient-recovery model and parameter estimation

The nutrient-recovery model was implemented on a two-dimensional grid and fitted independently to each colony. Experimental fluorescence image stacks were background-subtracted with a rolling-ball filter, cropped around each colony, Gaussian-smoothed, and block-averaged over 16 × 16-pixel regions for computational efficiency. Colony geometry was represented by an idealized circular mask whose time-dependent radius was obtained from a fit to the measured colony area expansion. Nutrient dynamics followed Equation 2, with consumption proportional to the colony growth activity, and fluorescence production was coupled to the nutrient field through the Hill response and radial height-weighting functions (Equations 4 and 5). Because diffusion is modeled in the plane of the pad, vertical transport through the agarose depth is absorbed into the effective parameter *κ* rather than represented explicitly.

Each colony was fitted to its two-dimensional fluorescence image stacks. For the triple constitutive reporter, the RFP and YFP channels were fitted jointly, and CFP was not fitted separately because it was assumed to share the YFP production response. For the single reporter repressilator, the mVenus and mCFP channels were fitted jointly, with the three repressor concentrations represented as latent model variables (Equation 6) and the repression Hill coefficient fixed at *m* = 4. Simulations were initialized from the experimental fluorescence fields at the first fitted frame, and each fit was started from a common initial parameter set for the corresponding construct.

Parameters were constrained to be positive and optimized in log-parameter space using bounded nonlinear least squares (scipy.optimize.least squares, Trust-Region Reflective). Residuals in each fluorescence channel were normalized by that channel’s mean observed intensity before being combined into the normalized mean squared error. The resulting best-fit parameters are phenomenological point estimates. The complete fitting code, parameter bounds, initial values, and per-colony fitted parameter sets are provided with the accompanying code and data.

### 4.7 Inter-colony diffusion model

Colony positions on each pad were extracted from the fluorescence images, and each colony was modeled as a point source of a diffusible factor with a common source strength *α* that was identical for all colonies. The concentration field evolved according to the two-dimensional diffusion equation (Equation 12), with no decay term, solved numerically on a grid matching the pad dimensions. The predicted arrival time at each colony was the time at which the local concentration first exceeded the threshold *q*_thresh_. Because the equation is linear in the source strength, *α* and *q*_thresh_ are not separately identifiable, so *α* was fixed to 1 and *q*_thresh_ is reported in simulation units. The two free parameters *D_q_* and *q*_thresh_ were estimated by bounded nonlinear least squares in log-parameter space (scipy.optimize.least squares), minimizing the squared differences between predicted and observed arrival times after subtracting their mean difference, so the fit tested relative differences in arrival time between colony positions rather than the absolute onset. A single global temporal offset, computed as the mean of observed minus simulated times, was added to the simulated arrival times for alignment. Agreement between model and data was quantified by Pearson’s *r* and the root mean square error.

### 4.8 Statistical analysis

Comparisons between dilution conditions used the Mann-Whitney *U* test. Correlations between Gompertz parameters and colony number were assessed by both Pearson and Spearman coefficients. Wave parameter comparisons used the Mann-Whitney test. Significance threshold was *p <* 0.05 throughout. All analyses were performed in Python (scipy.stats).

## 5 Data and code availability

All code will be deposited to GitHub before publication. Raw time-lapse data will be deposited to Zenodo before publication.

## 6 Acknowledgements

We thank Johan Paulsson for providing the single reporter repressilator (pLPT20) and triple reporter repressilator (pLPT107) plasmids (Addgene #85460 and #85525). G.Y.F. was supported by a School of Computing PhD studentship at Newcastle University. G.Y.F. was supported by UK Research and Innovation (UKRI) Biological Sciences Research Council (BBSRC) grant BB/Y008332/1.

## 7 Author information

### 7.1 Authors and Affiliations

**School of Computing, Newcastle University, Newcastle upon Tyne, NE1 7RU, United Kingdom**

Guillermo Yáñez Feliú, Timothy J. Rudge.

### 7.2 Contributions

G.Y.F. contributed to conceptual development, designed and performed experiments, developed image analysis tools, constructed mathematical models, performed data analysis, wrote the manuscript, and edited the manuscript. C.B. contributed to experimental data collection. T.J.R. supervised the project, contributed to conceptual development, designed the experiments, developed image analysis tools, constructed mathematical models, performed data analysis, wrote the manuscript, and edited the manuscript. All authors reviewed the final manuscript.

### 7.3 Corresponding author

Correspondence to G.Y.F. and T.J.R.

