## Supplementary material for "Self-generated environmental feedback drives traveling waves of gene expression in bacterial colonies"

### 1 Growth organization

#### 1.1 Growth-coupled repressilator model

An earlier modeling study predicted that coupling the repressilator to edge-dominated growth via protein dilution would generate traveling waves during active expansion [1]. We revisit this prediction using an updated radial velocity expression and show via simulation that this growth-coupled mechanism does not produce waves for a constitutive reporter ($\phi$ constant).

Mechanical interactions between cells and the substrate generate a characteristic growth-rate gradient in expanding *E. coli* microcolonies, with maximal growth at the colony edge and exponential decay toward the center:

$$\mu(r) = \mu_0 e^{-r/r_0}, \tag{S1}$$

where $r$ is the radial distance measured inward from the colony edge, $\mu_0$ is the maximal unconstrained growth rate, and $r_0$ is the characteristic decay length of the growth-rate profile. For biomass density $\rho$ moving with velocity field $\mathbf{v}$, conservation of biomass gives

$$\frac{\partial \rho}{\partial t} + \nabla \cdot (\rho \mathbf{v}) = \mu \rho. \tag{S2}$$

Assuming that the colony forms a densely packed monolayer with approximately constant biomass density, Equation S2 reduces to

$$\nabla \cdot \mathbf{v} = \mu. \tag{S3}$$

Let $s$ denote radial distance from the colony center, such that $r = R(t) - s$. Under radial symmetry, the absolute outward radial velocity $v_s(s,t)$ satisfies

$$\frac{1}{s}\frac{\partial}{\partial s}\left(s v_s\right) = \mu_0 e^{-[R(t)-s]/r_0}. \tag{S4}$$

Integrating from the colony center and imposing regularity at $s = 0$ gives

$$v_s(s,t) = \frac{\mu_0 r_0}{s}\left[(s - r_0)e^{-[R(t)-s]/r_0} + r_0 e^{-R(t)/r_0}\right]. \tag{S5}$$

Substituting $s = R - r$ gives the absolute outward radial velocity as a function of distance from the colony edge:

$$v_r(r,t) = \frac{\mu_0 r_0}{R - r}\left[(R - r - r_0)e^{-r/r_0} + r_0 e^{-R/r_0}\right]. \tag{S6}$$

The apparent singularity at the colony center is removable, with $v_r(R,t) = 0$ defined by continuity.

At the colony edge, $r = 0$ and $v_r(0,t) = v_{\text{front}} = dR/dt$, so

$$v_{\text{front}} = \frac{\mu_0 r_0}{R}\left[(R - r_0) + r_0 e^{-R/r_0}\right]. \tag{S7}$$

Solving Equation S7 for the maximal growth rate gives

$$\mu_0 = \frac{R\, v_{\text{front}}}{r_0\left[(R - r_0) + r_0 e^{-R/r_0}\right]}, \tag{S8}$$

and therefore

$$\mu_0 r_0 = \frac{R\, v_{\text{front}}}{(R - r_0) + r_0 e^{-R/r_0}}. \tag{S9}$$

Substituting Equation S9 into Equation S6 gives

$$
\begin{aligned}
v_r(r,t) &= \frac{1}{R-r}\left[\frac{R\,v_{\text{front}}}{(R-r_0)+r_0 e^{-R/r_0}}\right]\left[(R-r-r_0)e^{-r/r_0}+r_0 e^{-R/r_0}\right] \\
v_r(r,t) &= \frac{R\,v_{\text{front}}}{R-r}\,\frac{(R-r-r_0)e^{-r/r_0}+r_0 e^{-R/r_0}}{(R-r_0)+r_0 e^{-R/r_0}}.
\end{aligned}
\tag{S10}
$$

Figure S1A compares the normalized growth-rate profile, $\bar{\mu}(r)=\mu(r)/\mu_0=e^{-r/r_0}$, with the normalized outward radial velocity profile, $v_r(r)/v_{\text{front}}$, for a representative colony with $R/r_0=20$. The dotted line marks $r=r_0$, where the local growth rate has decreased to $1/e\approx 0.37$ of its value at the colony edge. Although the velocity at a given position depends on the cumulative growth occurring within the enclosed colony area, in the edge-dominated limit $r_0 \ll R$ the normalized velocity approaches

$$
\frac{v_r(r)}{v_{\text{front}}}\approx e^{-r/r_0}. \tag{S11}
$$

The near-overlap of the profiles is therefore an expected consequence of the large-colony limit and explains why the measured radial velocity field provides an indirect readout of the underlying spatial organization of growth.

Because $r$ is measured from the moving colony edge, the speed of a cell relative to that edge is

$$
u(r,t)=v_{\text{front}}(t)-v_r(r,t)\geq 0. \tag{S12}
$$

To derive the corresponding gene-expression equation, let $P_i(s,t)$ denote the concentration of protein $i$ in the fixed center coordinate. Its Eulerian material balance is

$$
\frac{\partial P_i}{\partial t}+v_s(s,t)\frac{\partial P_i}{\partial s}=\phi_i(\mathbf{P},t)-\left[\gamma_i+\mu(s,t)\right]P_i. \tag{S13}
$$

Defining $p_i(r,t)=P_i(R(t)-r,t)$ gives

$$
\frac{\partial p_i}{\partial t}=\frac{\partial P_i}{\partial t}+v_{\text{front}}\frac{\partial P_i}{\partial s},\qquad \frac{\partial p_i}{\partial r}=-\frac{\partial P_i}{\partial s}. \tag{S14}
$$

Using $v_s(R-r,t)=v_r(r,t)$ and substituting Equation S13 into Equation S14 gives

$$
\frac{\partial p_i}{\partial t}=\phi_i(\mathbf{p},t)-\left[\gamma_i+\mu(r,t)\right]p_i-\left[v_{\text{front}}(t)-v_r(r,t)\right]\frac{\partial p_i}{\partial r}. \tag{S15}
$$

Using the edge-relative speed defined in Equation S12, the transport equation becomes

$$\frac{\partial p_i}{\partial t} = \phi_i(\mathbf{p}, t) - [\gamma_i + \mu(r, t)]\, p_i - u(r, t)\frac{\partial p_i}{\partial r}. \tag{S16}$$

Here, $u \geq 0$ describes the rate at which a cell falls behind the advancing colony edge, corresponding to increasing distance $r$ from that edge. For a constitutive reporter, $\phi_i$ is constant. For the repressilator, $\phi_i$ depends on the concentration of the upstream repressor through a Hill repression function.

For simulation, we introduced the dimensionless variables

$$x = \frac{r}{r_0}, \qquad X = \frac{R}{r_0}, \qquad \tau = \mu_0 t, \qquad q_i = \frac{p_i}{K}, \tag{S17}$$

where $K$ is the common repression threshold of the repressilator. To simplify notation, the threshold-scaled protein concentrations $q_i$ are denoted by $p_i$ below. The remaining dimensionless quantities are

$$\bar{\mu} = \frac{\mu}{\mu_0}, \qquad \bar{\gamma}_i = \frac{\gamma_i}{\mu_0}, \qquad \bar{\phi}_i(\mathbf{q}, \tau) = \frac{\phi_i(K\mathbf{q}, \tau/\mu_0)}{\mu_0 K}, \qquad \bar{v}_r = \frac{v_r}{\mu_0 r_0}, \qquad \bar{u} = \frac{u}{\mu_0 r_0}. \tag{S18}$$

The dimensionless growth-rate profile is

$$\bar{\mu}(x) = e^{-x}. \tag{S19}$$

The dimensionless absolute radial velocity and front velocity are

$$\bar{v}_r(x, X) = \frac{(X - x - 1)e^{-x} + e^{-X}}{X - x}, \tag{S20}$$

and

$$\bar{v}_{\text{front}}(X) = \frac{(X - 1) + e^{-X}}{X}, \tag{S21}$$

respectively.

The dimensionless colony radius evolves according to

$$\frac{dX}{d\tau} = \bar{v}_{\text{front}}(X), \qquad X(0) = X_0. \tag{S22}$$

At the colony center, the corresponding limiting value is $\bar{v}_r(X, X) = 0$.

The dimensionless edge-relative speed is therefore

$$\bar{u}(x, X) = \bar{v}_{\text{front}}(X) - \bar{v}_r(x, X). \tag{S23}$$

For the repressilator, the dimensionless production functions were

$$\begin{aligned} \bar{\phi}_1(\mathbf{p}) &= \frac{\alpha}{1 + p_3^n}, \\ \bar{\phi}_2(\mathbf{p}) &= \frac{\alpha}{1 + p_1^n}, \\ \bar{\phi}_3(\mathbf{p}) &= \frac{\alpha}{1 + p_2^n}. \end{aligned} \tag{S24}$$

where $n$ is the Hill coefficient. For the constitutive reporter, the dimensionless production function was

$$\bar{\phi}_c = \alpha_c. \tag{S25}$$

The dimensionless maximal repressilator production rate $\alpha$ and constitutive production rate $\alpha_c$ are defined by

$$\alpha = \frac{\alpha_{\text{dim}}}{\mu_0 K}, \qquad \alpha_c = \frac{\alpha_{c,\text{dim}}}{\mu_0 K}, \tag{S26}$$

where $\alpha_{\text{dim}}$ and $\alpha_{c,\text{dim}}$ are the corresponding dimensional production rates.

The dimensionless gene-expression equation is

$$\frac{\partial p_i}{\partial \tau} = \bar{\phi}_i(\mathbf{p}, \tau) - [\bar{\gamma}_i + \bar{\mu}(x)]\, p_i - \bar{u}(x, X)\frac{\partial p_i}{\partial x}. \tag{S27}$$

The simulations in Fig. S1B,C used $X_0 = 1$, $\tau_{\text{max}} = 24$, $N = 400$, and $\Delta\tau = 0.002$. For the repressilator, $\alpha = 10^4$, $\bar{\gamma}_i = \bar{\gamma} = 1$ for $i = 1, 2, 3$, and $n = 2$, with initial conditions $p_1 = p_3 = 0$ and $p_2 = 5$. These parameter values were selected to reproduce the qualitative growth-coupled traveling-wave regime and were not fitted to the experimental data. For the constitutive reporter, $\alpha_c = 1$, $\bar{\gamma}_c = 1$, and $p_c(x, 0) = 0$. The value of $\alpha_c$ affects only the amplitude of the linear constitutive solution and not its normalized spatial profile.

At each time step, the exact dimensionless radial and front velocities were evaluated at the current colony radius, and the edge-relative speed $\bar{u}$ was used in Equation S27. Transport was discretized using a first-order upwind difference. At the moving colony edge, $x = 0$, where $\bar{u} = 0$, the transport term vanishes; the spatial derivative was set to zero in the numerical implementation. At the colony center, $x = X(\tau)$, the characteristic speed satisfies $\bar{u}(X, X) = dX/d\tau = \bar{v}_{\text{front}}$, so the center is a material boundary and no external protein-concentration boundary condition is required. Production and transport were evaluated explicitly, whereas linear degradation and growth dilution were treated semi-implicitly.

After updating the colony radius, the solution was interpolated onto a new uniform grid spanning $0 \leq x \leq X(\tau)$, with the previous center value assigned to grid points extending beyond the old domain. For RGB visualization, the three repressilator channels were independently contrast-normalized and rescaled by the largest channel at each pixel; color therefore represents relative oscillator phase rather than quantitative reporter amplitude. The simulated fields were subsequently mapped from distance from the moving edge, $x$, to radial position from the colony center, $s/r_0 = X - x$, to construct the kymographs.

Under this framework, the spatial growth-rate gradient causes cells at different distances from the colony edge to experience different rates of protein dilution. In the repressilator, this generates spatial phase differences that appear as traveling waves propagating from the expanding edge toward the colony center. A constitutive reporter has no oscillatory phase, and therefore forms a growth-dependent expression gradient without generating a traveling wave (Fig. S1B,C).

The model consequently predicts edge-dominated growth following Equations S1–S10, traveling waves in repressilator colonies during active expansion, and no traveling waves in constitutive reporter colonies.

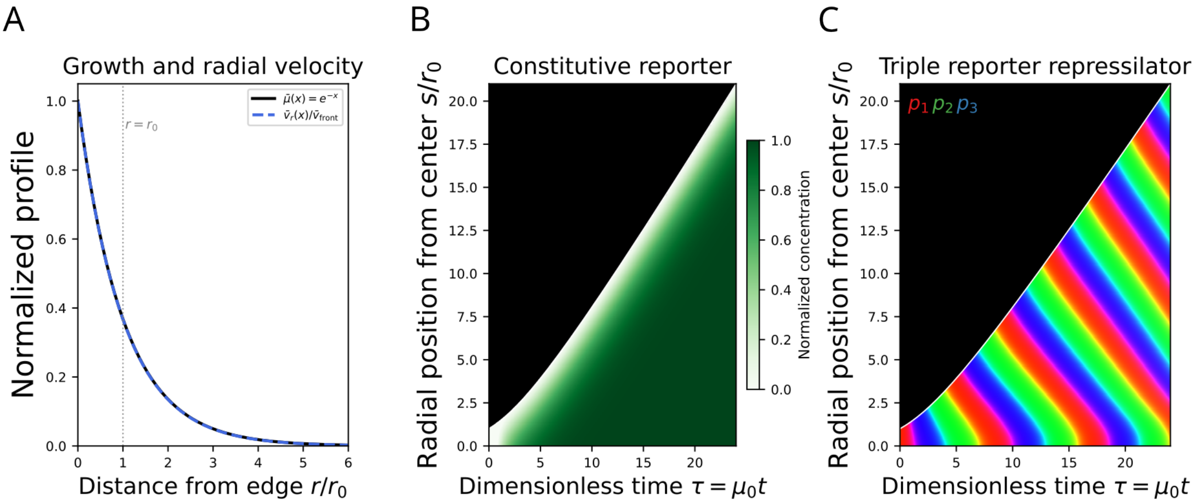


Figure S1: **Predictions of the growth-coupled repressilator model. (A)** Normalized growth-rate profile, $\bar{\mu}(r) = \mu(r)/\mu_0 = e^{-r/r_0}$, and normalized outward radial velocity, $v_r(r)/v_{\text{front}}$, for $R/r_0 = 20$. The dotted line marks $r/r_0 = 1$. **(B)** Simulated constitutive reporter. The reporter forms a smooth growth-dependent gradient that expands with the colony but does not generate a propagating wave. **(C)** Simulated triple-reporter repressilator. The independently normalized state variables $p_1$, $p_2$, and $p_3$, corresponding to mKate2, mVenus, and mCFP, are displayed in red, green, and blue, respectively. Color represents relative oscillator phase rather than quantitative reporter amplitude. Diagonal bands indicate phase waves propagating from the expanding colony edge toward the center. In **B** and **C**, the model was solved in the edge-relative coordinate $x = r/r_0$ and mapped to radial position from the colony center, $s/r_0 = X - x$, for visualization. White lines mark the colony boundary.

### 1.2 Velocity field computation and fitting

#### 1.2.1 Velocity computation

Radial velocity fields were computed from phase-contrast time-lapse images using mutual information image velocimetry, implemented via an ensemble grid approach. Briefly, each image was divided into overlapping windows of $64 \times 64$ pixels with a spacing of 32 pixels. For each window, the displacement between consecutive frames was estimated by maximizing the mutual information between the intensity distributions of the two frames, subject to a maximum displacement constraint of 19 pixels per frame. A colony mask was applied at each time point to restrict analysis to pixels within the growing colony. The resulting displacement field gives a velocity vector $(v_x, v_y)$ at each grid point and time point.

#### 1.2.2 Velocity processing

The raw displacement field was projected onto the radial direction at each grid point. The radial direction at each point was defined as the negative gradient of the Euclidean distance transform

(EDT) of the colony mask, pointing from the colony interior toward the edge. The radial velocity component was computed as the dot product of the displacement vector with this unit radial direction:

$$v_r = v_x \hat{g}_x + v_y \hat{g}_y, \tag{S28}$$

where $(\hat{g}_x, \hat{g}_y)$ is the unit outward radial vector at each grid point. A rigid-body drift correction was applied by subtracting the mean velocity across all grid points at each time point. The radial position of each grid point was taken as the mean EDT value within its window. Velocities were spatially smoothed by averaging each grid point with its $3 \times 3$ neighbors.

#### 1.2.3 Model fitting

For each colony, the colony front velocity $\hat{v}_{\mathrm{front}}(t) = dR/dt$ and colony radius $R(t)$ were estimated independently from the colony boundary at each time point. The single free parameter $r_0$, the characteristic decay length of the radial growth profile, was then estimated by minimizing the sum of squared residuals between the measured radial velocities and the model prediction (Equation S10), simultaneously across all time points and spatial positions within the fitting window ($t_0$ to $t_f$):

$$x^* = \arg\min_x \sum_{t,i,j} \left[\hat{v}_{\mathrm{front}}(t)\, \tilde{v}_r(r_{ij},\, R(t),\, r_0) - \bar{v}_{ij}(t)\right]^2, \tag{S29}$$

where $\tilde{v}_r = v_r / v_{\mathrm{front}}$ is the normalized model velocity profile, $r_{ij}$ is the mean radial position of grid window $(i, j)$, and $\bar{v}_{ij}(t)$ is the measured radial velocity at that window and time point. Minimization was performed using the Trust Region Reflective algorithm with $r_0$ parameterized as $r_0 = e^{x^*}$ to enforce positivity. This procedure yields a single time-averaged $r_0$ value per colony. The model was validated by comparing predicted and measured velocities pooled across all $n = 152$ colonies (8 excluded as quality-control failures, see below. $\mathrm{R}^2 = 0.892$; Fig. 1A). Colonies for which the fitted $r_0$ was less than or equal to 1 pixel were excluded as indicative of a failed fit (n=8 of 160, 5.0%). The model showed a small systematic overestimate of radial velocity (mean residual $-0.18$ pixels/frame, std 0.56 pixels/frame), consistent with the model assuming purely radial expansion while the velocimetry captures a small tangential noise component.

### 1.3 Justification of the $R > 200$ µm threshold for edge-dominated growth analysis

We restricted the characterization of the decay length $r_0$ to colonies with final radius $R > 200$ µm based on complementary analytical and empirical evidence. The analytical behavior of the velocity-profile model explains why $r_0$ becomes difficult to estimate in small colonies, while the clustering of failed fits and the dependence of per-colony fit quality on colony radius provide an empirical basis for the selected threshold.

#### 1.3.1 Analytical behavior in the small-colony limit

The radial velocity profile (Equation 1) contains the denominator

$$D(R, r_0) = (R - r_0) + r_0 e^{-R/r_0}. \tag{S30}$$

Taylor-expanding $e^{-R/r_0}$ for $R/r_0 \ll 1$ gives

$$D(R, r_0) = \frac{R^2}{2r_0} + \mathcal{O}\left(\frac{R^3}{r_0^2}\right), \tag{S31}$$

or, equivalently,

$$\frac{D(R, r_0)}{r_0} \approx \frac{1}{2}\left(\frac{R}{r_0}\right)^2. \tag{S32}$$

Although the complete velocity expression remains finite in this limit, its spatial profile approaches uniform radial expansion and becomes weakly sensitive to $r_0$. The decay length is therefore poorly identifiable from velocity data when the colony radius is small relative to $r_0$. Using the mean fitted decay length $\bar{r}_0 = 45.2$ µm across quality-control-passed colonies, $R = 200$ µm corresponds to $R/\bar{r}_0 \approx 4.4$ and $D/\bar{r}_0 \approx 3.4$.

#### 1.3.2 Empirical fit quality

Of the 160 colonies analyzed, eight returned fitted values $r_0 \leq 1$ pixel and were classified as failed fits. Their radii ranged from 106 to 210 µm, concentrating the failures at the lower end of the observed size range (Supplementary Fig. S2B). For the 152 quality-control-passed colonies, per-colony $R^2$ was computed from model-predicted and measured radial velocities using grid windows for which both velocities exceeded 0.1 pixels/frame. Fit quality was positively associated with colony radius (Pearson $r = 0.456$, $p < 0.0001$; Supplementary Fig. S2C). Colonies with $R \leq 200$ µm had a median per-colony $R^2$ of 0.11, compared with 0.85 for colonies with $R > 200$ µm (one-sided Mann–Whitney $U = 127$, $p < 0.0001$; Supplementary Fig. S2D). Several small colonies had negative $R^2$ values, indicating that the velocity-profile model performed worse than a constant mean predictor. Fitted decay lengths were also lower in the smaller-colony group. Colonies with $R \leq 200$ µm ($n = 13$) had $r_0 = 23.7 \pm 5.4$ µm, whereas colonies with $R > 200$ µm ($n = 139$) had $r_0 = 47.2 \pm 10.9$ µm (one-sided Mann–Whitney $U = 72$, $p = 2.2 \times 10^{-8}$, rank-biserial $r = 0.920$; Kolmogorov–Smirnov $D = 0.880$, $p = 6.4 \times 10^{-11}$). This difference is consistent with the distinct behavior of the fitted parameter in the small-colony and edge-dominated regimes. Together, the weak identifiability of $r_0$ in the small-colony limit, the clustering of failed fits at small radii, and the marked improvement in per-colony fit quality support the use of $R > 200$ µm as a conservative empirical threshold for characterizing edge-dominated growth.

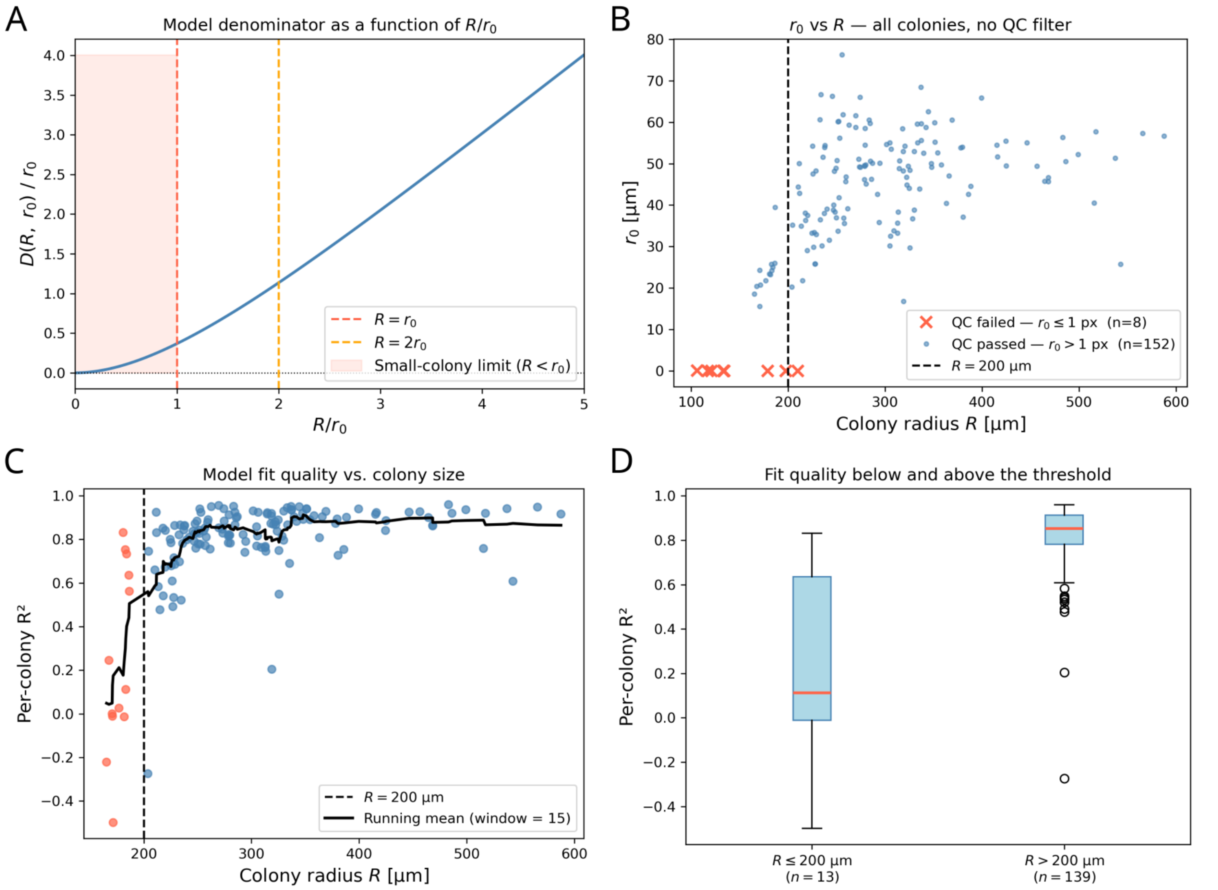


Figure S2: **Analytical and empirical justification of the $R > 200$ μm threshold. (A)** Normalized denominator of the radial velocity expression as a function of $R/r_0$. The denominator approaches zero quadratically in the small-colony limit, where the velocity profile approaches uniform expansion and becomes weakly sensitive to $r_0$. Dashed lines mark $R = r_0$ and $R = 2r_0$. **(B)** Fitted $r_0$ as a function of colony radius for all 160 colonies before quality-control filtering. Red crosses indicate failed fits with $r_0 \leq 1$ pixel ($n = 8$), and blue points indicate quality-control-passed fits ($n = 152$). The vertical dashed line marks $R = 200$ μm. **(C)** Per-colony $R^2$ as a function of colony radius. Points are colored by group: $R \leq 200$ μm (red, $n = 13$) and $R > 200$ μm (blue, $n = 139$). The solid line shows the running mean over 15 colonies ordered by radius. **(D)** Per-colony $R^2$ distributions below and above the 200 μm threshold. Median $R^2$ increased from 0.11 for $R \leq 200$ μm ($n = 13$) to 0.85 for $R > 200$ μm ($n = 139$, one-sided Mann–Whitney $U = 127$, $p < 0.0001$).

#### 1.3.3 Dependence of the fitted decay length on colony size

Across all quality-control-passed colonies, $r_0$ was positively associated with colony radius ($n = 152$, Pearson $r = 0.403$, $p < 0.0001$; Supplementary Fig. S3A). A weaker positive association persisted after restricting the analysis to colonies in the edge-dominated regime ($R > 200$ μm, $n = 139$, Pearson $r = 0.233$, $p = 0.0058$; Supplementary Fig. S3B). Smaller colonies therefore strengthen the overall $r_0$–$R$ association but do not account for it entirely. Within the edge-dominated regime, $r_0$ retains a weak positive dependence on colony radius. Because this dependence is substantially slower than the increase in $R$, it is consistent with the observed decrease in $r_0/R$ with colony radius (Fig. 1B). Because a single time-averaged $r_0$ was fitted for each colony, this cross-colony analysis does not resolve how the decay length changes during the growth of an individual colony.

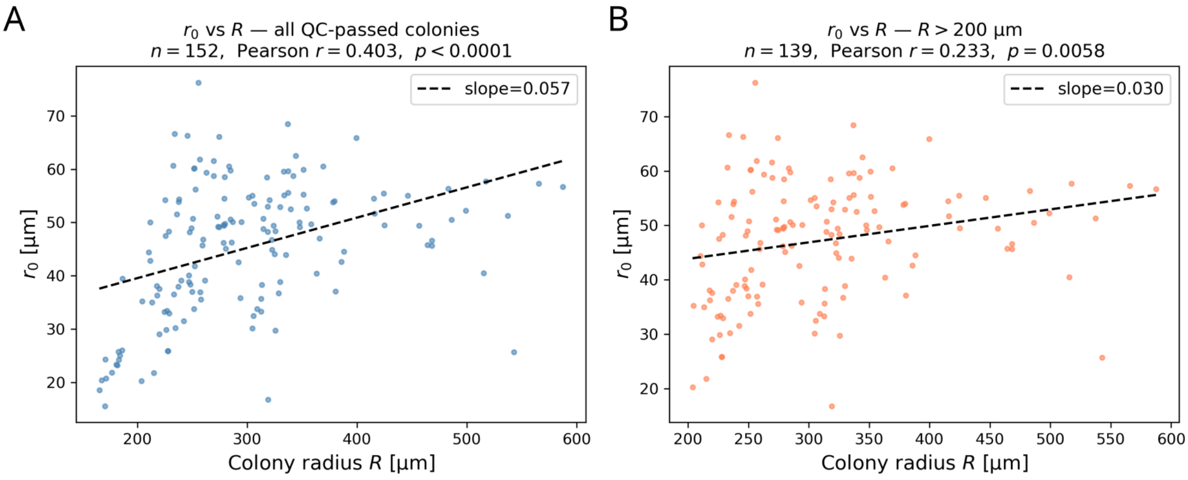


Figure S3: **Dependence of the fitted decay length on colony size.** **(A)** Fitted decay length $r_0$ as a function of colony radius $R$ for all quality-control-passed colonies ($n = 152$, Pearson $r = 0.403$, $p < 0.0001$). **(B)** The same relationship after restricting the analysis to colonies in the edge-dominated regime ($R > 200$ μm, $n = 139$, Pearson $r = 0.233$, $p = 0.0058$). Dashed lines show ordinary least-squares linear fits.

### 1.4 Gompertz growth characterization

We fitted the area expansion of each colony over time to a Gompertz model, which yields the asymptotic final area $A_\infty$, the maximum growth rate $\mu_m$, the lag phase $\lambda$, the time of maximum growth $t_m$, and the growth stop time $t_s$. All five parameters were negatively associated with the number of colonies per pad $N$ (Table S1). Thus, colonies in denser environments were smaller, had lower maximum growth rates, and reached growth termination earlier.

The growth stop time captured this relationship most directly. Colonies under high-density seeding stopped growing earlier than those under low-density seeding (Fig. S4A), and $t_s$ declined continuously with $N$ across both seeding conditions rather than separating into two discrete groups (Fig. S4B). This continuous association is consistent with growth termination responding gradually to the shared growth environment rather than being determined by an intrinsic colony timer.

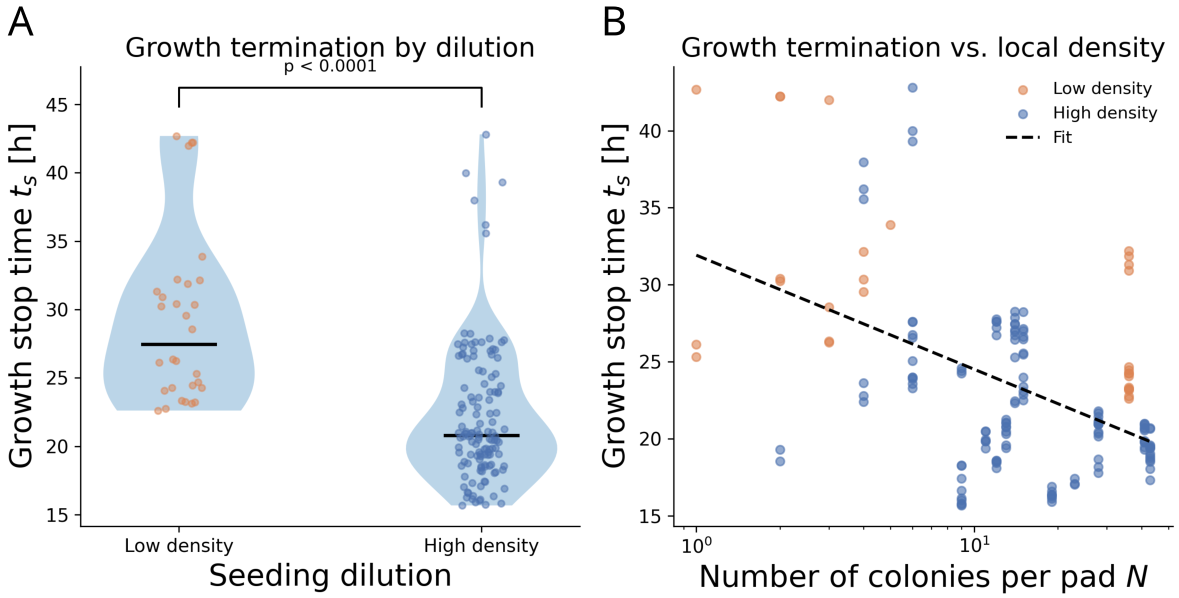


Figure S4: **Growth termination varies with colony density. (A)** Growth stop time $t_s$ under low- and high-density seeding conditions. Violins show the distributions, points show individual colonies, and horizontal lines indicate medians. Colonies under high-density seeding stopped growing significantly earlier than those under low-density seeding (median $t_s = 20.8$ h versus 27.5 h; Mann–Whitney $p < 0.0001$). **(B)** Growth stop time $t_s$ per colony as a function of the number of colonies per pad $N$. Points are colored by seeding condition. The dashed line is a linear fit against $\log_{10} N$ (slope $= -7.40$ h per decade). Growth stop time was negatively associated with colony density across both seeding conditions (Pearson $r = -0.381$, $p = 6.6 \times 10^{-7}$), consistent with a graded response to the shared growth environment rather than an intrinsic colony timer.

| Parameter | Pearson $r$ | $p$ (Pearson) | Spearman $\rho$ | $p$ (Spearman) |
|---|---|---|---|---|
| $A_\infty$ | $-0.436$ | $8.1 \times 10^{-9}$ | $-0.476$ | $1.9 \times 10^{-10}$ |
| $\mu_m$ | $-0.384$ | $5.3 \times 10^{-7}$ | $-0.414$ | $5.1 \times 10^{-8}$ |
| $\lambda$ | $-0.345$ | $8.0 \times 10^{-6}$ | $-0.282$ | $3.0 \times 10^{-4}$ |
| $t_m$ | $-0.349$ | $6.2 \times 10^{-6}$ | $-0.310$ | $6.8 \times 10^{-5}$ |
| $t_s$ | $-0.381$ | $6.6 \times 10^{-7}$ | $-0.376$ | $9.7 \times 10^{-7}$ |

Table S1: Correlation between the number of colonies per pad $N$ and Gompertz growth parameters ($n = 160$ colonies). All Pearson and Spearman $p$-values were below 0.001, indicating significant negative associations consistent with density-dependent growth dynamics.

### 1.5 Density-dependent growth

If colonies competed for a strictly fixed resource pool, the total colony area per pad would be constant regardless of how many colonies the pad contained. When the two seeding conditions were compared directly, total colony area per pad was not significantly different between them (Fig. S5A), which, considered alone, appeared consistent with a fixed pad-level carrying capacity.

Resolving total colony area against $N$ showed that this was not the case. On linear axes, a fixed pad-level carrying capacity would appear as a horizontal relationship. Instead, total colony area increased above the constant-area reference at high $N$ (Fig. S5B), which is inconsistent with total area remaining fixed. This complements the log-log scaling in Fig. 2C, where the same data fall between the fixed carrying-capacity and fixed individual-size limits.

Finally, the two seeding conditions occupied largely distinct regions of $(\log N, \log \bar{A})$ space and were separated by a linear support vector machine boundary with a cross-validated classification accuracy of 94% (Fig. S5C). This shows that the two inoculum conditions sampled different ranges of colony number and mean colony area. Across these ranges, higher colony numbers were associated with smaller individual colonies.

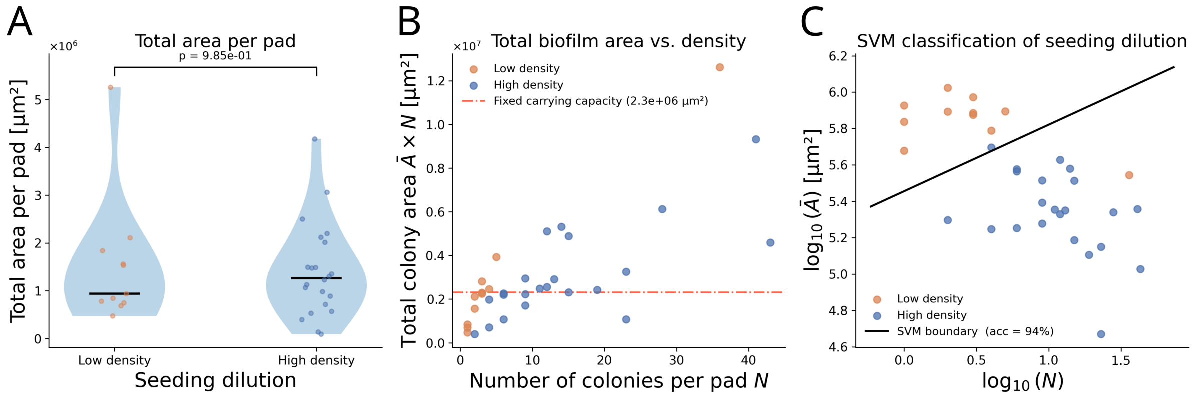


Figure S5: **Analysis of density-dependent growth. (A)** Total colony area per pad under low- and high-density seeding conditions. Violins show the distributions, points show individual pads, and horizontal lines indicate medians. Total colony area was not significantly different between conditions (Mann–Whitney $p = 0.985$), showing that fewer, larger colonies and more, smaller colonies produced comparable total colony areas. **(B)** Total colony area $\bar{A}N$ per pad as a function of colony number $N$, shown on linear axes. The red dash-dotted line indicates the median total colony area across all pads ($\bar{A}N = 2.3 \times 10^6$ µm$^2$), used as a constant-area reference. Total colony area exceeded this reference at high $N$, which is inconsistent with a fixed pad-level carrying capacity. **(C)** Linear support vector machine classification of seeding condition in $(\log N, \log \bar{A})$ space. Orange indicates low-density seeding and blue indicates high-density seeding ($n = 33$ pads). The solid line shows the SVM decision boundary. Leave-one-out cross-validated classification accuracy was 94%, showing that the two seeding conditions occupied largely distinct regions of $(N, \bar{A})$ space.

### 1.6 Scaling of colony area with colony number

We consider how mean colony area $\bar{A}$ and total biofilm area $\bar{A} \times N$ are expected to depend on the number of colonies per pad $N$ under two limiting hypotheses, and compare these with the observed scaling.

**Fixed carrying capacity.**

If each pad supports a fixed total biomass $K$, set by its finite and identical resource content, and this biomass is divided among $N$ colonies, then the mean colony area is

$$\bar{A} = \frac{K}{N} \propto N^{-1}, \tag{S33}$$

and the total biofilm area is constant,

$$\bar{A} \times N = K \propto N^{0}. \tag{S34}$$

Under this hypothesis, mean colony area falls as $1/N$ and total area is independent of $N$.

**Fixed individual size.**

If instead each colony grows to the same final size regardless of its neighbours, as in a scenario with infinite resources, then mean colony area is constant,

$$\bar{A} \propto N^{0}, \tag{S35}$$

and total biofilm area grows linearly with colony number,

$$\bar{A} \times N \propto N^{1}. \tag{S36}$$

**Observed scaling.**

Fitting power laws to the data gives

$$\bar{A} \propto N^{-0.47} \approx N^{-1/2}, \qquad \bar{A} \times N \propto N^{0.53} \approx N^{1/2}. \tag{S37}$$

Both exponents lie approximately halfway between the two limiting cases. Mean colony area decreases as $\approx 1/\sqrt{N}$ rather than $1/N$, and total biofilm area increases as $\approx \sqrt{N}$ rather than remaining constant or growing linearly. The two exponents are consistent with one another, since by definition the exponent of $\bar{A} \times N$ must equal one plus the exponent of $\bar{A}$ $(-0.47 + 1 = 0.53)$.

This intermediate scaling indicates partial competition: colony growth is suppressed by local crowding, but colonies do not partition a strictly fixed resource pool, consistent with competition acting locally rather than uniformly across the entire pad.

### 2 Gene expression organization

All plasmid diagrams were constructed using SBOLCanvas [2] in accordance with the SBOL Visual standard.

#### 2.1 Plasmids

The pAAA plasmid (figure S6) contains three constitutively expressed fluorescent reporters: RFP (mRFP1), YFP (EYFP), and CFP (ECFP) [3]. Each reporter is under the control of separate constitutive promoters (J23101), allowing simultaneous expression of all three fluorescent proteins. This plasmid was used to follow the expression dynamics of three constitutively expressed reporters simultaneously in bacterial colonies.

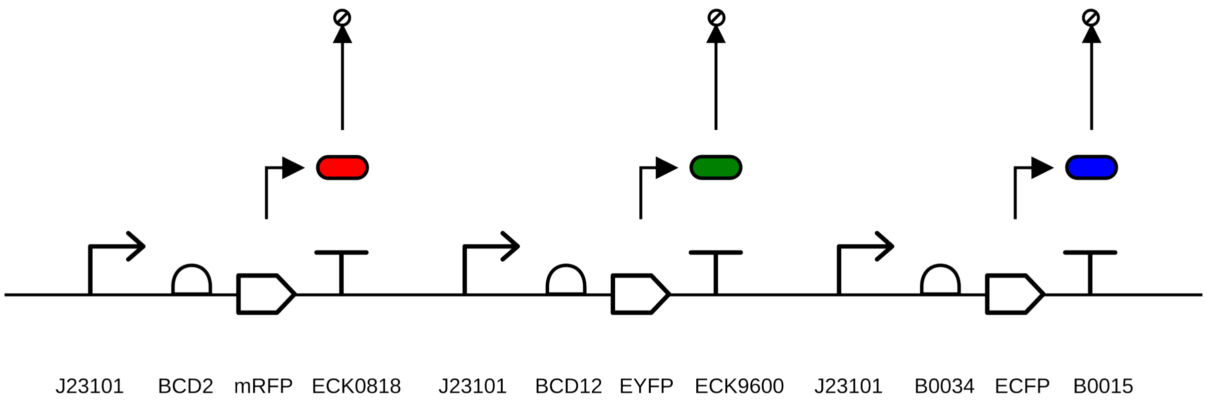


Figure S6: **pAAA SBOL plasmid map.** The pAAA plasmid contains three constitutively expressed fluorescent reporters: RFP (mRFP1), YFP (EYFP), and CFP (ECFP), each controlled by separate constitutive promoters (J23101).

The pLPT20 plasmid (figure S7) is a single reporter repressilator containing three repressors ($\lambda$cI, LacI, and TetR) arranged in a feedback loop to generate oscillatory gene expression dynamics [4]. The YFP reporter (mVenus) is regulated by the TetR repressor, while CFP (mCFP) is constitutively expressed and provides an internal reference channel. This configuration allows the regulated YFP channel and the constitutive CFP channel to be followed together in growing colonies.

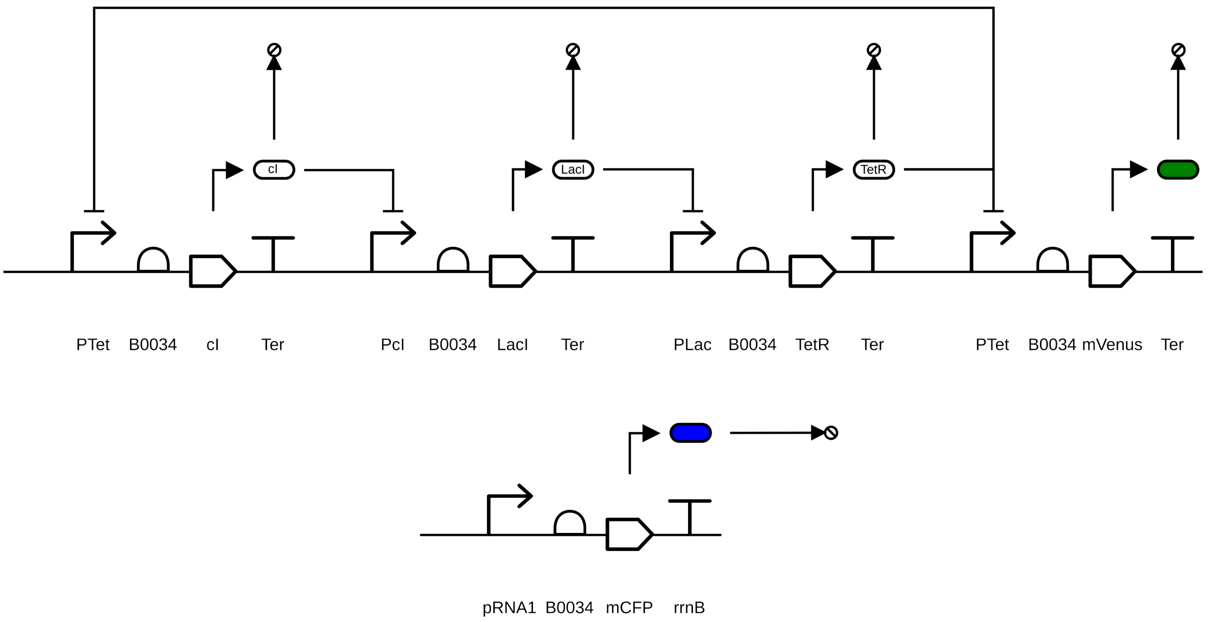


Figure S7: **Single reporter repressilator pLPT20 SBOL plasmid map.** The pLPT20 plasmid contains a single regulated reporter, YFP (mVenus), and a constitutively expressed CFP (mCFP). YFP expression is controlled by the TetR repressor, while CFP is expressed from a constitutive promoter and serves as an internal reference channel. This plasmid is used to follow regulated and constitutive reporter dynamics in bacterial colonies.

The pLPT107 plasmid (figure S8) is a triple reporter repressilator, with three fluorescent reporters — RFP, YFP, and CFP — each linked to a different repressor: LacI, $\lambda$cI and TetR, respectively. Unlike the single reporter **pLPT20**, this plasmid tracks the activity of all three repressors, allowing visualization of oscillatory gene expression dynamics across bacterial colonies.

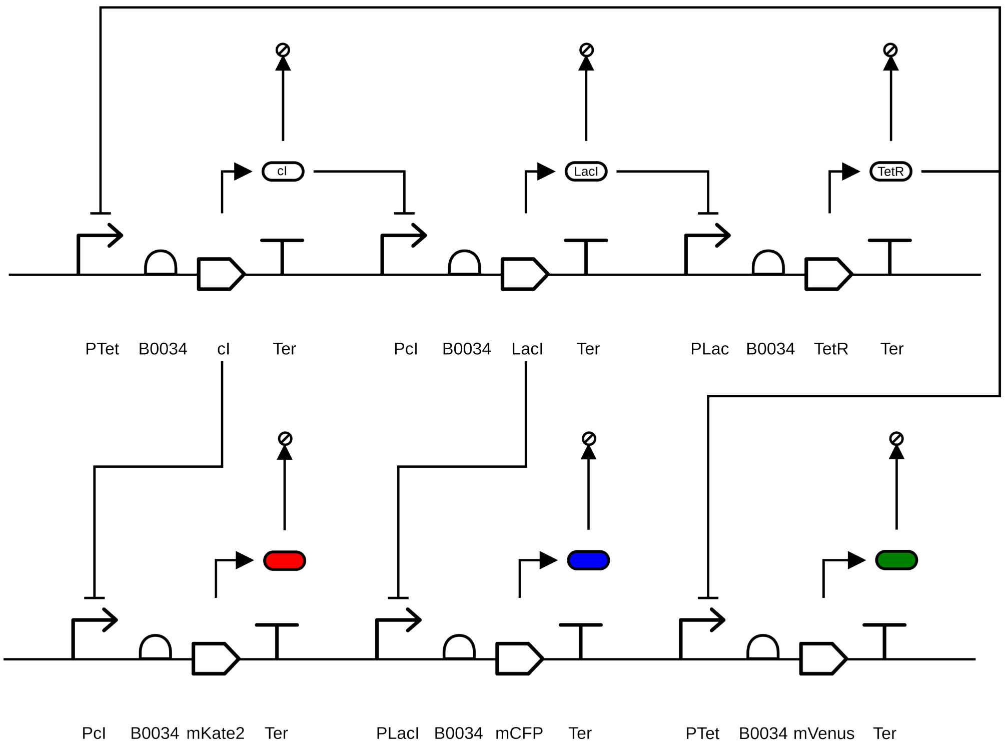


Figure S8: **Triple reporter repressilator pLPT107 SBOL plasmid map.** The pLPT107 plasmid contains three fluorescent reporters—RFP (mKate2), YFP (mVenus), and CFP (mCFP)—each linked to one of the three repressors: LacI, $\lambda$cI and TetR, respectively. These repressors regulate each other in a feedback loop, generating oscillatory gene expression dynamics. The fluorescent reporters allow real-time tracking of these dynamics across the colony, providing insight into the temporal regulation of gene expression.

### 2.2 A nutrient-recovery model recapitulates intra-colony traveling waves after growth arrest

The nutrient-recovery model was fitted independently to each of several colonies per construct. Normalized mean squared error was calculated by dividing the residuals in each fluorescence channel by the mean observed intensity of that channel and averaging the squared normalized residuals across all finite spatial and temporal data points. For the triple constitutive reporter, seven colonies from two experiments were fitted (three from 2023-11-28 and four from 2023-11-30). For the single reporter repressilator, nine colonies from one experiment (2023-12-08) were fitted, of which one (position 13) was excluded because its fitted LacI repression constant was implausible ($K_{\mathrm{LacI}} \approx 1551$, compared with a retained-fit median of approximately 0.076) and its normalized mean squared error was the highest of the set. Position 2 was retained because it showed good model-data agreement despite extreme values in some latent repressor parameters, consistent with parameter compensation and weak identifiability rather than a failed fit. Positions 15 and 12 were selected as the representative colonies shown in Fig. 6 and in the main-text parameter tables; both also had the lowest normalized mean squared error within their respective datasets. Per-colony normalized mean squared errors are reported in Table S2, and the representative values are placed alongside the across-colony medians and interquartile ranges in Tables S3 and S4. These interquartile ranges describe variation among best-fit point estimates from different colonies and should not be interpreted as confidence intervals or parameter-identifiability bounds.

| Experiment | Position | Normalized MSE | Status |
|---|---|---|---|
| *Triple constitutive reporter* ($n = 7$) | | | |
| 2023-11-28 | 0 | 0.0369 | Included |
| 2023-11-28 | 4 | 0.0963 | Included |
| 2023-11-28 | 8 | 0.1209 | Included |
| 2023-11-30 | 14 | 0.0786 | Included |
| 2023-11-30 | 15 | 0.0170 | Included (representative) |
| 2023-11-30 | 28 | 0.0229 | Included |
| 2023-11-30 | 33 | 0.0247 | Included |
| *Single reporter repressilator* ($n = 9$, one excluded) | | | |
| 2023-12-08 | 0 | 0.0280 | Included |
| 2023-12-08 | 1 | 0.0215 | Included |
| 2023-12-08 | 2 | 0.0187 | Included |
| 2023-12-08 | 4 | 0.0418 | Included |
| 2023-12-08 | 6 | 0.0350 | Included |
| 2023-12-08 | 7 | 0.0391 | Included |
| 2023-12-08 | 11 | 0.0209 | Included |
| 2023-12-08 | 12 | 0.0179 | Included (representative) |
| 2023-12-08 | 13 | 0.0791 | Excluded |

Table S2: Per-colony normalized mean squared error for the nutrient-recovery model fits. Positions 15 and 12 were selected as the representative colonies shown in Fig. 6 and also had the lowest normalized mean squared error within their respective constructs. Position 13 of the single reporter repressilator was excluded because it combined the highest normalized mean squared error with an extreme fitted $K_{\mathrm{LacI}}$ value.

| Parameter | Units | Rep. (pos15) | 2023-11-28 | 2023-11-30 | Pooled median [IQR] |
|---|---|---|---|---|---|
| $\kappa$ | $m^2\ s^{-1}$ | $1.33 \times 10^{-13}$ | $1.27 \times 10^{-13}$ | $1.32 \times 10^{-13}$ | $1.30 \times 10^{-13}$ [$1.27 \times 10^{-13}$, $1.32 \times 10^{-13}$] |
| $\nu_u$ | – | 9.35 | 5.47 | 9.33 | 9.30 [5.48, 9.33] |
| $\gamma_r$ | $s^{-1}$ | $3.15 \times 10^{-4}$ | $2.71 \times 10^{-4}$ | $3.25 \times 10^{-4}$ | $2.85 \times 10^{-4}$ [$2.72 \times 10^{-4}$, $3.25 \times 10^{-4}$] |
| $\gamma_y$ | $s^{-1}$ | $7.34 \times 10^{-6}$ | $5.88 \times 10^{-6}$ | $7.34 \times 10^{-6}$ | $7.32 \times 10^{-6}$ [$5.88 \times 10^{-6}$, $7.34 \times 10^{-6}$] |
| $\alpha_r$ | a.u. $s^{-1}$ | 9.91 | 7.86 | 9.40 | 7.97 [7.81, 9.40] |
| $\alpha_y{}^{\dagger}$ | a.u. $s^{-1}$ | $7.74 \times 10^{-6}$ | $7.74 \times 10^{-6}$ | $7.74 \times 10^{-6}$ | $7.74 \times 10^{-6}$ |
| $\beta_r$ | a.u. $s^{-1}$ | 0.534 | 0.703 | 0.533 | 0.538 [0.533, 0.703] |
| $\beta_y$ | a.u. $s^{-1}$ | $1.70 \times 10^{-3}$ | $1.82 \times 10^{-3}$ | $1.70 \times 10^{-3}$ | $1.70 \times 10^{-3}$ [$1.70 \times 10^{-3}$, $1.82 \times 10^{-3}$] |
| $\tau_r$ | – | 0.430 | 0.361 | 0.432 | 0.430 [0.363, 0.432] |
| $\tau_y{}^{\dagger}$ | – | 2.52 | 2.52 | 2.52 | 2.52 |
| $n$ | – | 22.2 | 18.8 | 22.1 | 22.0 [18.8, 22.1] |
| $\sigma_h$ | – | 0.300 | 0.522 | 0.291 | 0.314 [0.291, 0.518] |

Table S3: Triple constitutive reporter parameters across the seven fitted colonies. Rep.: representative colony (position 15). The 2023-11-28 and 2023-11-30 columns show the experiment-specific medians for $n = 3$ and $n = 4$ colonies, respectively. Within-experiment variation was small for most parameters, while differences between experiment medians contributed to the pooled interquartile ranges, most clearly for $\sigma_h$ and $\nu_u$. The pooled summaries therefore combine between-colony and between-experiment variation. $^{\dagger}$The parameters $\alpha_y$ and $\tau_y$ remained unchanged from their initialized values in every fit and were therefore not independently informed by the optimization; their absence of spread should not be interpreted as precise estimation.

| Parameter | Units | Rep. (pos12) | Median [IQR] ($n = 8$) |
|---|---|---|---|
| $\kappa$ | $m^2\ s^{-1}$ | $2.07 \times 10^{-13}$ | $1.64 \times 10^{-13}$ [$1.34 \times 10^{-13}$, $1.80 \times 10^{-13}$] |
| $\nu_u$ | dimensionless | 1.27 | 2.15 [1.70, 3.54] |
| $\sigma_h$ | dimensionless | 0.602 | 0.897 [0.587, 1.34] |
| $K_{\mathrm{LacI}}$ | a.u. | 0.0759 | 0.0752 [0.0607, 0.0909] |
| $K_{\mathrm{TetR}}$ | a.u. | 50.5 | 55.2 [50.1, 58.5] |
| $K_{\mathrm{cI}}$ | a.u. | 2.89 | 20.1 [5.06, 37.7] |
| $\gamma_{\mathrm{repr}}$ | $s^{-1}$ | $3.87 \times 10^{-5}$ | $3.24 \times 10^{-5}$ [$1.44 \times 10^{-5}$, $4.26 \times 10^{-5}$] |
| $\gamma_y$ | $s^{-1}$ | $6.36 \times 10^{-6}$ | $5.84 \times 10^{-6}$ [$3.91 \times 10^{-6}$, $8.03 \times 10^{-6}$] |
| $\gamma_c$ | $s^{-1}$ | $2.55 \times 10^{-4}$ | $2.18 \times 10^{-4}$ [$1.60 \times 10^{-4}$, $3.21 \times 10^{-4}$] |
| $\alpha_y$ | a.u. $s^{-1}$ | 0.561 | 0.602 [0.554, 0.665] |
| $\alpha_c$ | a.u. $s^{-1}$ | 6.20 | 5.28 [4.82, 7.82] |
| $\beta_{\mathrm{LacI}}$ | a.u. $s^{-1}$ | $2.35 \times 10^{-6}$ | $5.51 \times 10^{-6}$ [$3.52 \times 10^{-6}$, $6.78 \times 10^{-5}$] |
| $\beta_{\mathrm{TetR}}$ | a.u. $s^{-1}$ | $2.59 \times 10^{-3}$ | $2.37 \times 10^{-3}$ [$2.00 \times 10^{-3}$, $2.95 \times 10^{-3}$] |
| $\beta_{\mathrm{cI}}$ | a.u. $s^{-1}$ | 0.0255 | 0.0314 [0.0249, 0.0358] |
| $\beta_c$ | a.u. $s^{-1}$ | 2.05 | 2.05 [1.84, 2.96] |
| $\tau_{\mathrm{LacI}}$ | dimensionless | 0.817 | 0.779 [0.730, 0.846] |
| $\tau_{\mathrm{TetR}}$ | dimensionless | 0.766 | 0.728 [0.685, 0.766] |
| $\tau_{\mathrm{cI}}$ | dimensionless | 0.519 | 0.415 [0.374, 0.470] |
| $\tau_c$ | dimensionless | 0.474 | 0.324 [0.295, 0.409] |
| $n$ | dimensionless | 25.4 | 15.7 [13.4, 21.1] |
| $R_{\mathrm{LacI}}(0)$ | a.u. | 0.0845 | 0.0536 [0.0256, 0.0663] |
| $R_{\mathrm{TetR}}(0)$ | a.u. | $2.43 \times 10^{-3}$ | $2.81 \times 10^{-3}$ [$2.40 \times 10^{-3}$, $3.58 \times 10^{-3}$] |
| $R_{\mathrm{cI}}(0)$ | a.u. | $1.69 \times 10^{-3}$ | $5.87 \times 10^{-4}$ [$5.15 \times 10^{-4}$, $1.05 \times 10^{-3}$] |

Table S4: Single reporter repressilator parameters across the eight retained colonies, excluding position 13. Rep.: representative colony (position 12). The wide interquartile ranges of some repression constants, particularly $K_{\mathrm{cI}}$, and initial repressor concentrations are consistent with parameter compensation and weak identifiability among the latent repressilator variables, while also including biological variation between colonies. The representative parameter set should therefore be interpreted as one solution that reproduces the observed reporter dynamics rather than as a uniquely determined mechanistic parameterization.

### 2.3 An inter-colony suppression wave suggests diffusive coupling

Figure 7B summarizes the arrival time of the suppression wave as a distribution per pad, which shows that arrival times vary within a pad and stay comparable across pads, but it does not preserve the position of each colony. To test whether this variation is spatially organized, we plotted the arrival time of each colony at its measured position within the pad (Figure S9). Within pads with several colonies, neighboring colonies tend to share similar arrival times, and arrival times vary gradually from one side of the pad to the other rather than appearing at random positions.

This spatial structure is the direct observation that motivates the diffusion model in the main text. If the suppression wave reflected a uniform change in the shared medium or an intrinsic cellular timer triggered at the same developmental stage, arrival times would not depend on colony position. The graded spatial pattern is instead consistent with a factor produced by the colonies and spreading through the shared medium, reaching each colony at a time set by its position relative to the sources. This figure therefore provides the spatial evidence behind the staggered arrival times in Figure 7B and the diffusion model fit in Figure 7C.

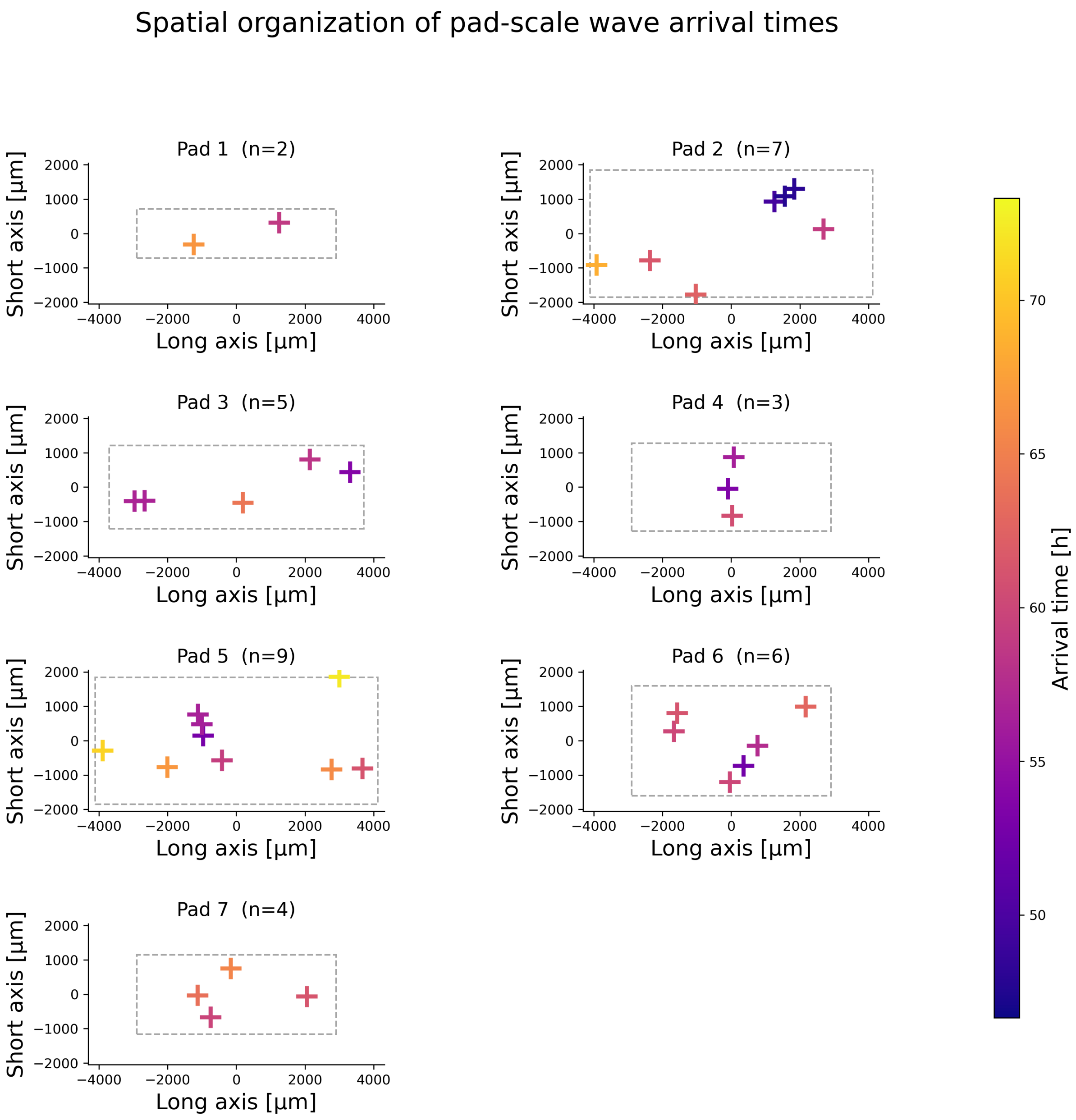


Figure S9: **Spatial organization of pad-scale wave arrival times.** Each panel shows the spatial position of colonies within one pad, with points colored by the experimentally measured arrival time of the pad-scale suppression wave. Grey dashed rectangles indicate the imaged pad region. Arrival times vary across colonies within the same pad, showing that the second wave is not globally synchronized but emerges at different times depending on colony position within the shared environment.
